# Hippocampal Theta Phase Precession Supports Memory Formation and Retrieval of Naturalistic Experience in Humans

**DOI:** 10.1101/2023.06.05.543539

**Authors:** Jie Zheng, Mar Yebra, Andrea G.P. Schjetnan, Clayton Mosher, Suneil K. Kalia, Jeffrey M. Chung, Chrystal M. Reed, Taufik A. Valiante, Adam N. Mamelak, Gabriel Kreiman, Ueli Rutishauser

## Abstract

Linking different experiences together is a key aspect of episodic memory. A potential neural mechanism for linking sequential events over time is phase precession, which causes neurons to fire progressively earlier in time relative to theta-frequency local field potential oscillations. However, no direct link between phase precession and behaviorally assessed memory encoding or retrieval success has been established. We recorded the activity of single neurons and local field potentials in the human medial temporal lobe (MTL) while participants encoded and retrieved memories of movie clips. Transient brief theta bouts and theta phase precession were observed following cognitive boundaries during movie watching as well as following stimulus onset during memory retrieval. Phase precession was dynamic, with different neurons exhibiting phase precession in different task periods. The strength of phase precession provided information about memory encoding and retrieval success that was not available in firing rates, thereby linking the temporal code established by phase precession to behaviorally assessed memory strength. These data reveal phase precession during non spatial memory in humans and provide direct neural evidence for a functional role of phase precession in episodic memory.

## Introduction

From significant life events to mundane everyday moments, episodic memory plays a crucial role in shaping our perception of the world and our sense of self. Our brains not only store snapshots of individual events but also temporally weave together sequential contiguous moments into rich and coherent memories for future use. A fundamental open question in human memory is, how do we encode and retrieve memories of continuous experience?

Theoretical work suggests that the encoding, binding, and compressing of sequential events into a coherent memory might rely on a temporal neural code^1–4^. A key motivation for these theories is that the precise timing at which spikes of hippocampal neurons occur relative to ongoing field potential activity in the theta frequency range carries information about both the content of a given experience as well as the location of a given event within a sequence^5–9^. A key mechanism that gives rise to temporal coding is phase precession, whereby neurons spike in progressively earlier phases of ongoing theta oscillations relative to the onset of either a significant event^6,10^ or the location of the animal in the case of place cells^11–13^. As a result of phase precession, the phase of spiking relative to theta is indicative of the elapsed time since the onset of the event. Phase precession was first discovered in rodents^11^ and has since been investigated extensively in the context of spatial coding of self^12^ and others^14^. Phase precession has since also been reported in other species, including marmosets^15^, bats^16^ and humans^17, 18^. A key insight of the work in marmosets, bats, and humans is that phase precession is robustly present even if the underlying theta activity is only present intermittently in bouts^16^. Recent work further generalizes the presence of phase precession towards non-spatial domains, demonstrating its relevance to auditory stimuli identity encoding^6, 19^, odor identity encoding^6^, time in an event sequence^20–22^, and sleep^20^.

Despite its prevalence, the precise functional role that phase precession plays in memory remains elusive. Phase precession coordinates hippocampal cell assemblies such that different cells fire in order within a single theta cycle^13, 23^ in a manner that is conductive to synaptic plasticity^2, 24^. Recent rodent^6^ and human studies^17, 25^ report that phase precession occurs most strongly following event onsets or event transitions. Based on these data, it has been hypothesized that a function of phase precession is to organize neural activity such that a cohesive memory can be formed and later be retrieved. However, so far no direct link between the extent of phase-precession and episodic memory-related behavior has been established.

We examined whether phase precession strength is related to whether a non spatial episodic memory is encoded or retrieved in humans during continuous semirealistic experience. Our results make three key contributions. First, we report non spatial phase precession in humans after event boundaries during naturalistic experience. Second, phase precession also occurred during recognition memory and order memory retrieval. Third, the strength of phase precession reflected participants’ memory encoding and retrieval success. Overall, our findings extend phase precession to non-spatial episodic memory and demonstrate a direct link between phase precession and memory behaviors in humans.

## Results

### Task and neural recording

22 subjects with drug-resistant epilepsy participated in the experiment (3 new participants in addition to the 19 for which we already published single-neuron but not field potential data^26^; Supplementary Table S1). The task consisted of three parts: encoding, scene recognition, and time discrimination (see *Methods*). During encoding, participants watched 90 movie clips of approximately 8s length each. Each clip was novel and consisted of either a single continuous shot (Fig. 1b, referred to as “no boundary clips”) or several different shots separated by event boundaries (Fig. 1a, “boundary clips”). After watching all 90 clips once, participants performed two memory tests. During the scene recognition test (Fig. 1c), participants indicated whether the presented frame was “old” (from watched clips) or “new” (not watched) via button press. During the time discrimination test (Fig. 1d), order memory was evaluated by asking participants to indicate which one of two frames shown on the screen appeared earlier in time in the movie clips just watched. Patients performed well in both memory tests, with accuracy of 73% ± 10% and 70% ± 12% for scene recognition and time discrimination test, respectively (scene recognition: *p* < 4x10^-4^; time discrimination: *p* < 3x10^-3^; one sample *t*-test with respect to chance level). Participants were implanted with depth electrodes for clinical evaluation while performing the task (see participant demographics in Supplementary Table 1 and spike quality metrics in Supplementary Fig. 1).

**Fig. 1.**
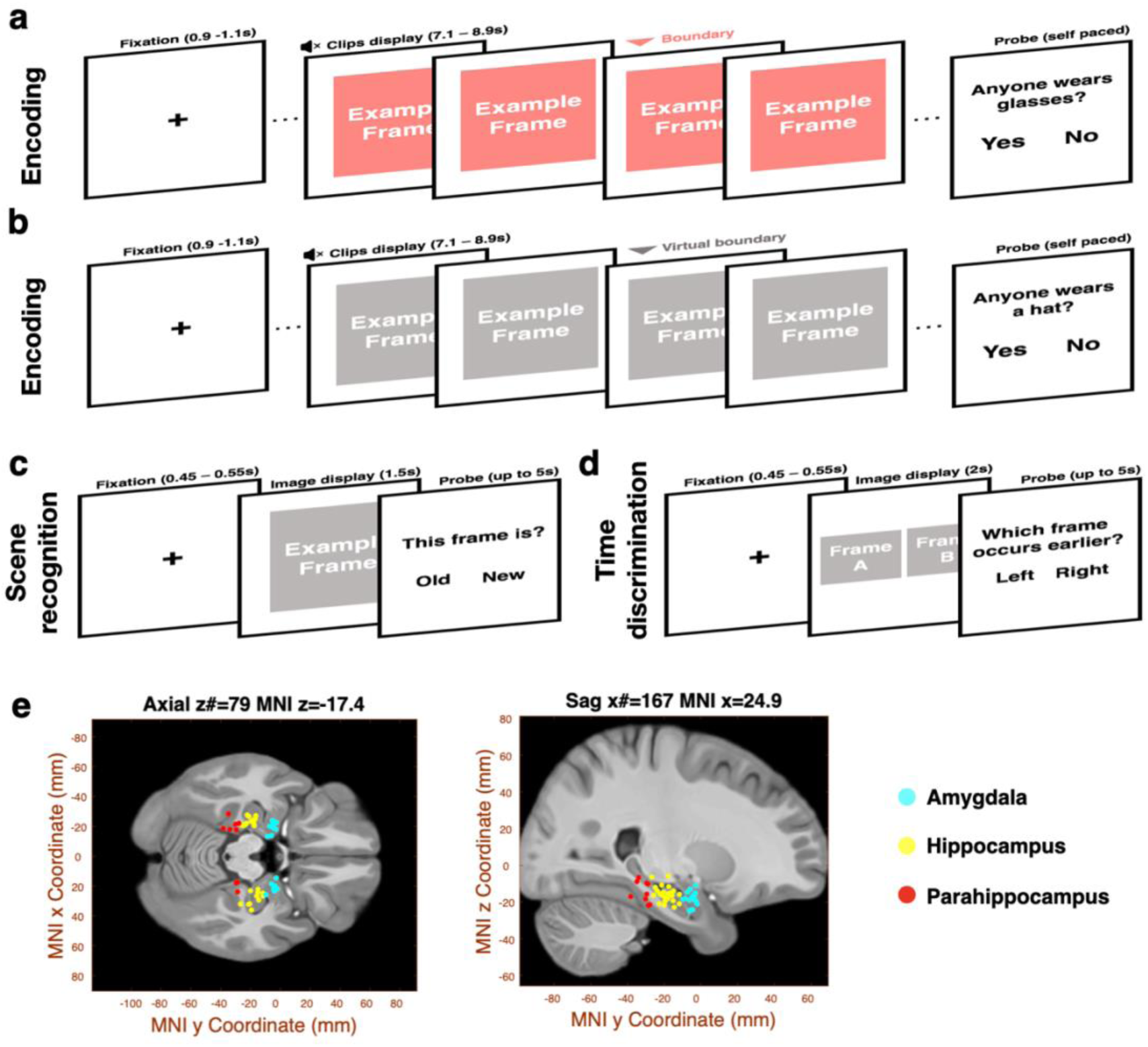
Experiment and electrode locations. **(a-c)** Schematics of the three task stages. **(a** and **b)** Encoding. Participants watched a series of silent clips and were instructed to answer yes/no questions that appeared randomly after every four to eight clips^26^. These clips either contained cuts to scenes from the same or different movie (a, boundary clips) or no cuts (b, no boundary clips). The red triangle in (a) marks the timepoint of the boundary in an example boundary clip. The gray triangle in (b) refers to the timepoint of 4 second in an example no boundary clip. **(c)** Scene recognition. Participants were presented with a static image and were asked to indicate whether the image was “old” (shown in the watched clips) or “new”. **(d)** Time discrimination. Participants were presented with two images side by side and were asked to indicate whether the left or right frame appeared first in the watched clips. See^26^ for more detailed information about the task. Due to copyright issues, all original images have been removed but are available upon reasonable request. **(e)** Illustration of the location of the 50 microelectrodes in the medial temporal lobe across 22 participants (see participants’ demographics in Supplementary Table 1) that are included in this study. Shown is a slice from a template brain CIT168 (see *Methods*), with microelectrodes plotted as individual dots, and color-coded for different brain regions (Amygdala: cyan; Hippocampus: yellow; Parahippocampal gyrus: red). Montreal Neurological Institute (MNI) coordinates for all microelectrodes in the plot are listed in Supplementary Table 2.

We quantified phase precession using simultaneously recorded single-neuron activity and local field potentials (LFP) from microwire electrodes implanted in the hippocampus, amygdala, and parahippocampal gyrus (Fig. 1e and Supplementary Table 2), jointly referred to as the MTL. We also recorded data from three frontal areas, including orbitofrontal cortex (OFC), anterior cingulate (ACC), and pre-supplementary motor area (preSMA), which we analyzed to compare the prevalence of phase precession between the frontal lobe (in total 41 microwires, 433 neurons) and MTL (in total 50 microwires, 503 neurons).

### Transient theta bouts in the MTL are prevalent following cognitive boundaries

As the internal reference “clock” for neuronal spiking^8^, we first assessed the properties of theta-band local field potentials (LFPs) recorded from the MTL (see electrode locations in Fig. 1e and electrode Montreal Neurological Institute Coordinates in Supplementary Table 2). Recordings from the amygdala, hippocampus, and parahippocampal gyrus revealed large non-rhythmic fluctuations in the LFPs (Fig. 2a, 2c, 2e). Averaging power across the entire task, the LFPs recorded in these regions had approximately 1/*f*-shaped power spectra, with no apparent peaks in the conventional theta range of 4-8Hz (Fig. 2a-f, Supplementary Fig. 2a-c and 2e-g).

**Fig. 2.**
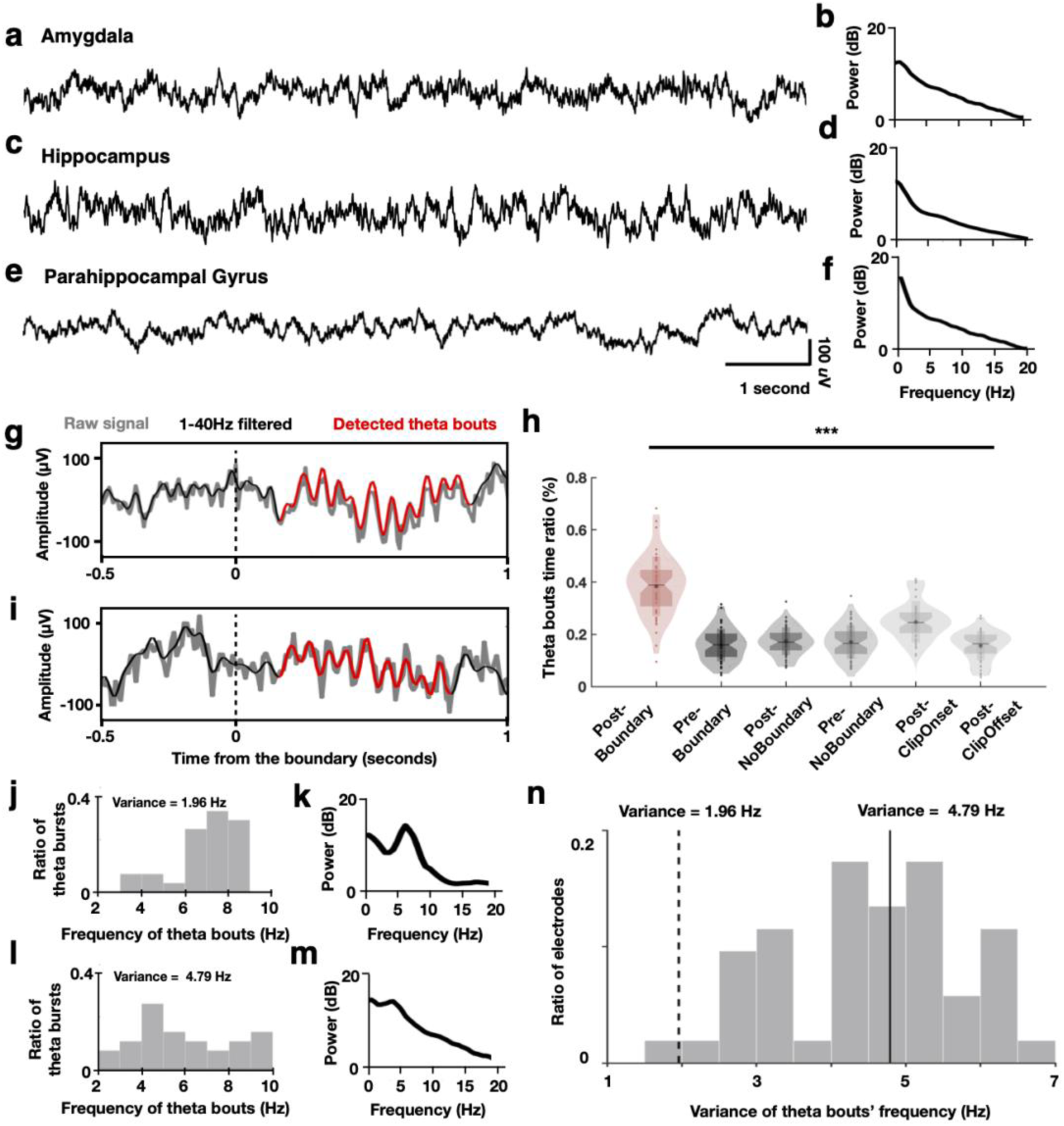
Characteristics of theta bouts during encoding. **(a-f)** Example local field potentials (LFPs) recorded from microelectrodes located in the amygdala (a), hippocampus (c), and parahippocampal gyrus (e). (b,d,f) LFP power spectra from the same example electrodes in a, c, e recorded throughout the entire task. **(g** and **i)** Examples of detected theta bouts. Shown is the raw LFP (gray), 1-40Hz bandpass filtered LFP (black), and detected theta bout (red). t=0 is the timepoint when a boundary occurs. (**h**) Proportion of time occupied by theta bouts within different 1-second time windows (analysis windows) before (Pre Boundary) and after boundaries (Post-Boundary) in boundary clips, before (Pre-NoBoundary) and after (Post-NoBoundary) the midpoint in no boundary clips, after clip onsets (Post-ClipOnset) and clip offsets (Post-ClipOffset) of all the clips. Each dot represents one microelectrode. The shaded violin shape represents the data distribution with lower end of 1^st^ percentile and top end of 99^th^ percentile. The top edge and bottom edge of the shaded rectangle represents the mean + std. and mean – std., respectively. \*\*\**p* < 0.001 (ANOVA test across all the analysis windows). **(j** and **l)** Frequency of all detected theta bouts within the Post-Boundary time windows for the two example microelectrodes shown in (g) and (i), respectively. The inset shows the variance. **(k** and **m)** Power spectra of the LFPs across all the Post Boundary windows from the microelectrodes in (g) and (i), respectively. **(n)** Variance of the frequency of theta bouts across all recorded microwires in the MTL. Solid and dashed lines mark the variance of the two example microelectrodes shown in (g) and (i), respectively.

We next searched for brief ‘theta bouts’ in the time domain on a cycle-by-cycle basis^27^ and quantified each oscillatory cycle by its amplitude, period and waveform symmetry (see *Methods*). This analysis revealed transient “theta bouts” (see examples highlighted in Fig. 2g and 2i; see *Methods*), consistent with previous findings in humans^27–29^ and other far-sensing species^15, 16^. On average, 14% ± 4% of the entire recording time was occupied by theta bouts. The likelihood of theta bout occurrence varied across different task periods. During encoding, considering 1s long periods, theta bouts were more likely to occur following cognitive boundaries in boundary clips (Fig. 2h; *F*_5,244_ = 12.4, *p* < 6x10^-7^; one-tailed ANOVA), with theta bouts occuping 36.4% ± 8.7% of post-boundary time windows. In contrast, theta bouts only occupied 16.2% ± 5.9%, 18.3% ± 4.7%, 16.8% ± 6.7%, 25.2% ± 8.8% and 15.6% ± 4.9% of the 1s long time periods before cognitive boundaries in boundary clips, before and after no boundaries in no boundary clips, after clip onsets and offsets in all clips, respectively. The frequency of theta bouts was heterogeneous, spanning 2Hz to 10Hz. This was true even when comparing different bouts detected on the same microelectrode (Fig. 2j and 2l shows two example microelectrodes). The variance of theta bout frequency changed considerably across electrodes, with some electrodes having theta bouts with relatively consistent frequencies (Fig. 2l, mostly from 6Hz to 8Hz). As a result, the microelectrodes with relatively low variance in bout-by-bout frequency changes had power spectra with prominent theta peaks when only considering the 1s period following cognitive boundaries (Fig. 2k), whereas those with larger variance had at best a modest peak (Fig. 2m). Across all electrodes, low variance was relatively rare, with most electrodes exhibiting relatively large variance in the frequency of theta bouts (Fig. 2n; variance of theta bouts frequency for all the electrodes: 4.18 ± 1.14 Hz).

Consistent with the findings during encoding, theta oscillations during memory retrieval were also relatively rare, resulting in approximately 1/*f* -shaped power spectra (Supplementary Fig. 2a-c and 3e-g). The incidence of theta bouts increased following image onset in both retrieval tasks relative to baseline (Supplementary Fig. 2d and 2h; % of time; scene recognition: 22.2% ± 6.3% vs. 12.7% ± 3.8%; *F*_2,97_ = 6.88, *p* < 2x10^-3^; time discrimination: 21.6% ± 4.2% vs. 14.2% ± 3.7%*F*_2,97_ = 7.11, *p* < 2x10^-3^; one-tailed ANOVA). Theta bout prevalence was also relatively low during the later memory retrieval periods (Supplementary Fig. 2d and h). Compared to encoding, the total time occupied by theta bouts was substantially larger during encoding relative to that following image onsets in both memory retrieval tasks (See Supplementary Fig. 2i; encoding *vs* scene recognition: 36.4% ± 8.7% *vs.* 21.3% ± 4.2%, *p* < 1x10^-5^, Kolmogorov-Smirnov test; encoding *vs* time discrimination: 36.4% ± 8.7% *vs.* 22.4% ± 6.2%, *p* < 2x10^-7^, Kolmogorov-Smirnov test). Lastly, the frequency of detected theta bouts following image onsets varied more compared to theta bouts following cognitive boundaries during encoding (Supplementary Fig. 2j; scene recognition *vs* encoding: 5.25 ± 0.98 Hz *vs* 4.18 ± 1.14 Hz, *P* < 2x10^-3^, Kolmogorov-Smirnov test; time discrimination *vs* encoding: 5.47 ± 0.93 Hz *vs* 4.18 ± 1.14 Hz, *P* < 1x10^-4^, Kolmogorov Smirnov test). In sum, these data show that LFPs in the human MTL contain prominent but relatively rare and short transient bouts of activity in the theta range. Theta bouts were most common following the occurrence of cognitive boundaries during encoding.

### Theta phase precession of individual neurons mostly occurs following boundaries

Next, we examined whether the spiking of single neurons exhibited theta phase precession during memory formation and retrieval of continuous experience. As theta bouts occurred predominantly after boundaries during encoding (Fig. 2h), we started by assessing phase precession within the post boundary windows in boundary clips. Given the wide range of theta bout frequencies (Fig. 2n), we quantified phase precession with respect to activity within a range of theta frequencies between 2 to 10Hz. Similar to the methods used in rodents^11, 13, 30^, we quantified phase precession by computing the circular correlation^31^ between the theta phase of spikes and elapsed time after boundaries within each microwire. We assessed the significance of phase precession using a shuffle-based permutation procedure (see *Methods*). We quantified elapsed time as the total accumulated theta phase rather than absolute time, a metric we refer to as ‘unwrapped phase’ throughout. This is a common method to assess phase precession first introduced by Mizuseki et al^32^ and widely used in the context of nonstationary LFPs^16, 17^. For example, 0° in the x-axis in Fig. 3a marks the time of boundary onset and 1080° marks the time after three cycles of theta have elapsed.

**Fig. 3.**
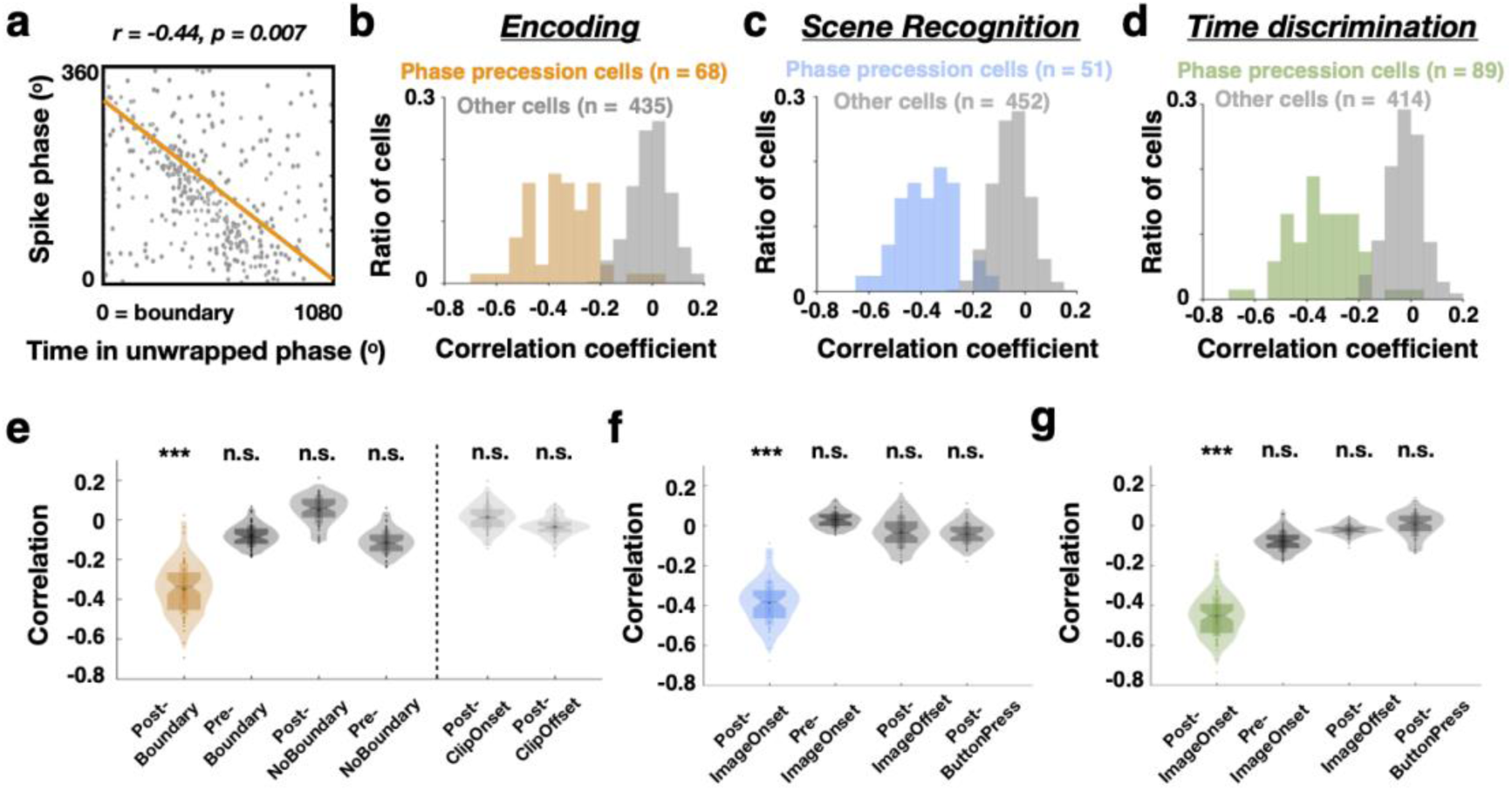
Prevalence of theta phase precession across different task stages. **(a)** Example hippocampal phase precession neuron (detected from the microelectrode shown in Fig. 2g). The spiking of this neuron exhibited phase precession after boundaries during encoding. The spiking phases relative to local theta (y-axis) are plotted as a function of time in unwrapped theta phase (i.e., how many theta cycles have passed, Methods). Each dot shows one spike within three theta cycles (1080 degrees on the x-axis) relative to boundaries in boundary clips. The orange line indicates the fitted correlation between neuronal spiking phase and time in unwrapped theta phase, with its correlation value (*r*) and significance level (*p*) listed on top of the plot. (**b** - **d**) Distribution of correlation coefficients for all MTL neurons demonstrating significant phase precession during encoding (b, orange), scene recognition (c, blue) or time discrimination (d, green). Distribution of correlation coefficients for neurons without significant phase precession is plotted in gray. Note that strong phase precession is indicated by negative correlation coefficients. (**e**) Comparison of phase precession strength during encoding for phase precession neurons in (b). Stronger phase precession is indicated by more negative correlations. Plotted are circular-linear correlation coefficients computed using spikes fired within three theta cycles before (Pre-Boundary) and after (Post-Boundary) boundaries in boundary clips, after clip onsets (Post-ClipOnset) and after clip offsets (Post-ClipOffset) in boundary clips, also before (Pre-NoBoundary) and after (Post-NoBoundary) the midpoint in no boundary clips. (**f** and **g**) Comparison of phase precession strength during scene recognition (f) or time discrimination (g) for phase precession neurons in (c) and (d), respectively. Plotted are circular-linear correlation coefficients computed using spikes fired within three theta cycles before (Pre-ImageOnset) and after image onsets (Post-ImageOnset), after image offsets (Post-ImageOffset) and after making a memory choice (Post-buttonPress). In (e-g), each dot represents one neuron. The shaded violin shape represents the data distribution with lower end of 1^st^ percentile and top end of 99^th^ percentile. The top edge and bottom edge of the shaded rectangle represent the mean ± std., respectively. \*\*\**p*< 0.001 (one-sample *t*-test against zero), n.s. = not significant.

We found that 68/503 (13.5%; above chance, *p* < 7x10^-4^, permutation test) of neurons in the MTL demonstrated significant phase precession following cognitive boundaries during encoding as assessed using this method (*p* < 0.05, permutation test; see example in Fig. 3a and *Methods*). Fig. 3b shows the distribution of correlation coefficients for the selected 68 neurons. As time elapsed following cognitive boundaries, most phase precession neurons (66/68, 97%) spiked progressively earlier relative to ongoing theta (from 360° to 0°), characterized by a negative spike-phase correlation (Fig. 3b). Phase precession neurons exhibited an average correlation coefficient of -0.38 ± 0.11 (*p* < 3 x 10^-8^, one-sample *t*-test), which is comparable to the strength of phase precession reported previously (Feng et al, 2015; Terada et al, 2017; Schmidt et al, 2009). By definition, phase precession neurons had more negative (*p* < 3x10^-4^, Kolmogorov-Smirnov test) correlation coefficients compared to non-phase precession neurons (−0.07 ± 0.13, mean ± std; *p* = 0.26, one-sample *t*-test). To evaluate the robustness of phase precession, we also computed phase precession using two alternative methods, which demonstrated consistent findings (Supplementary Fig. 3 and Supplementary Table 3; see *Methods*).

Was phase precession specific to the 1s period of time following cognitive boundaries? For the neurons showing phase precession after cognitive boundaries (n = 68, Fig 3b orange), phase precession strength was not significantly different from chance for other periods of time during encoding (Fig. 3e). These findings suggest that phase precession is present during memory formation of continuous experience following cognitive boundaries.

### Phase precession is task dependent and anatomically specific

We next turned to assessing phase precession during memory retrieval. We found that 10.1% (51/503) and 17.7% (89/503) of MTL neurons showed phase precession (*p* < 0.05, permutation test) when participants were presented with tested images during scene recognition (Fig. 3c) and time discrimination (Fig. 3d), respectively. These neurons exhibited an average correlation coefficient of -0.41 ± 0.09 (*p* < 2 x 10^-8^, one-sample *t*-test) during scene recognition and -0.40 ± 0.12 during time discrimination. By definition, these correlation coefficients were more negative than those of non-phase precession neurons (scene recognition: *p* < 2x10^-4^; time discrimination: *p* < 2x10^-4^; Kolmogorov-Smirnov test). Similar to theta bouts, the observed phase precession was prominent following the onsets of image display but not during baseline, following image offset, and during the button press period (Fig. 3f and 3g).

We found that phase precession is a dynamic process. Among all the phase precession neurons observed in the MTL across encoding, scene recognition, and time discrimination (n = 117), half (61/117, 47.9%) demonstrated phase precession exclusively for only one task stage (Fig. 4d; Fig. 4a-c shows an example). Of the neurons that only showed phase precession in one task, the majority did so during time discrimination (40/61, 65.6%) rather than encoding (18/61, 29.5%) or scene recognition (3/61, 4.9%). We also observed neurons that showed phase precession during multiple task stages (Fig. 4d; 56/117, 52.1%), with most of these neurons showing phase precession for all three task stages (35/56, 62.5%). The strength of phase precession varied across different task stages (see example in Fig. 4a-c), with more negative correlations (indicating stronger phase precession) during the two memory retrieval tasks compared to encoding (Fig. 4e and 4f, scene recognition – encoding: *r_diff* = - 0.19 ± 0.09, *p* < 3x10^-4^; time discrimination – encoding: *r_diff* = -0.24 ± 0.11, *p* < 6x10^- 7^; one-sample *t*-test). Comparing the two retrieval tasks, phase precession was stronger during time discrimination compared to scene recognition (time discrimination – scene recognition: *r_diff* = -0.11 ± 0.04, *p* < 0.03, one sample *t*-test). The observed differences in phase precession strength could not be explained by differences in firing rates because neurons had comparable firing rates across the three tasks (Supplementary Fig. 4a: scene recognition - encoding: *Fr_diff =* -0.18 ± 0.41 spikes/s*, p* = 0.33, one-sample *t*-test; time discrimination - encoding: *Fr_diff =* -0.14 ± 0.38 spikes/s, *p* = 0.24, one sample *t*-test). Lastly, we asked whether phase precession strength varied as a function of different types of trials within a given task. Phase precession during scene recognition was significantly stronger following onset of images that were old compared to images that were novel (−0.26 ± 0.14 vs. -0.11 ± 0.07, p < 4 x 10^-3^, paired *t*-test; see Supplementary Fig. 5), suggesting a role of precession in retrieval. Together, these results indicate that the strength by which a given neuron exhibits phase precession is a dynamic process that varies both as a function of the task and trial type within the task.

**Figure 4.**
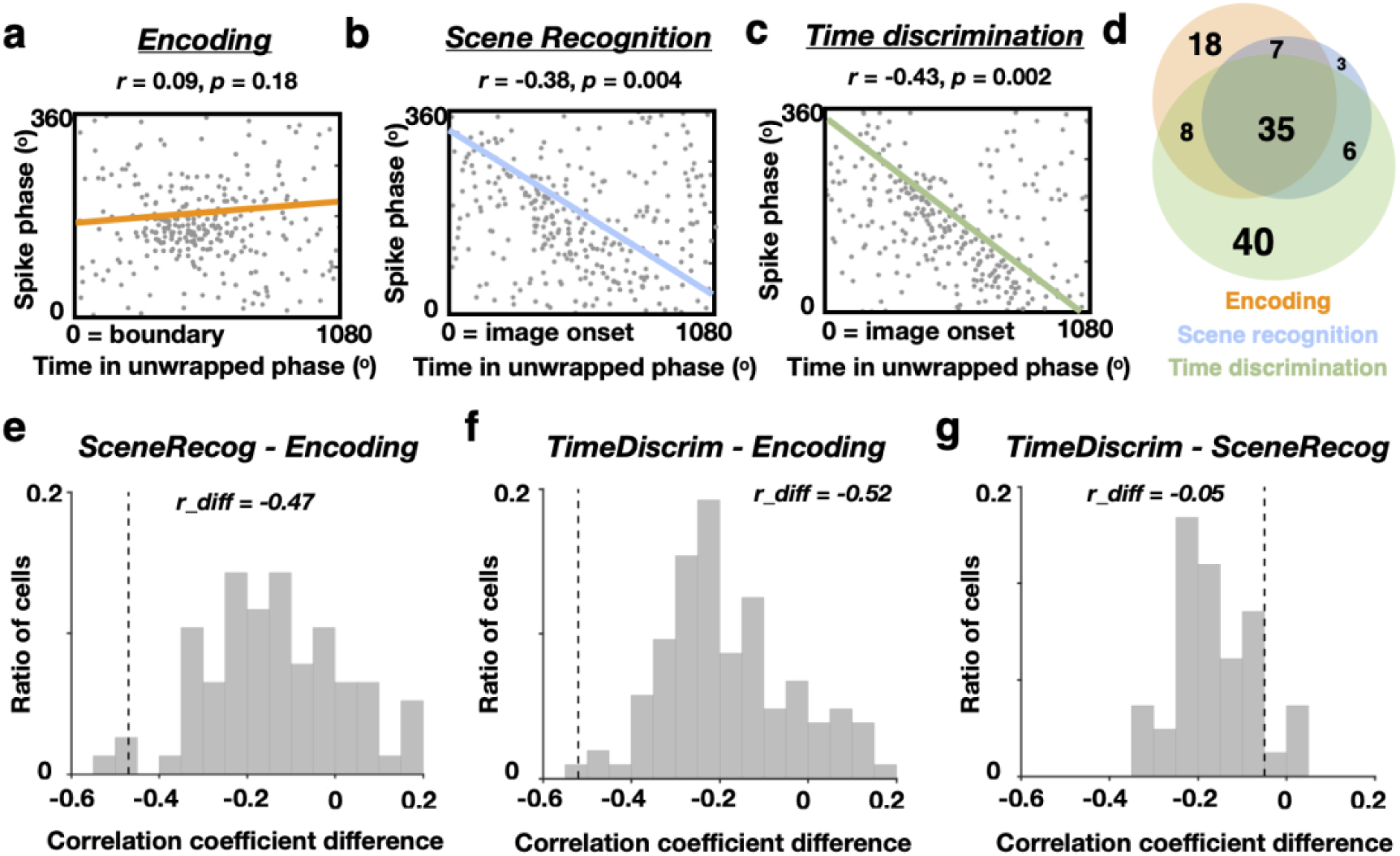
Comparison of phase precession strength across different task stages. (**a**-**c**) Example hippocampal neuron whose spiking exhibited phase precession during recognition and time discrimination, but not encoding. Shown are spike phases as a function of time in unwrapped theta phase, displayed separately for encoding (a; t=0 is boundary in boundary clips), scene recognition (b; t=0 is image onset) and time discrimination (c; t=0 is image onset). Colored lines indicate the fitted correlation between spike phase and time in unwrapped theta phase. The correlation value (*r*) and its statistical significance (*p*) are listed on top of each plot. (**d**) Number of MTL neurons showing significant phase precession during encoding (orange), scene recognition (blue), time discrimination (green), and combinations thereof. (**e**) Difference in phase precession strength between encoding and scene recognition for neurons that show phase precession for encoding and/or scene recognition (i.e., orange plus blue circles in panel d). (**f**) Difference in phase precession strength between encoding and time discrimination for neurons that showed significant phase precession during encoding and/or time discrimination (i.e., orange plus green circles in d). (**g**) Difference in phase precession strength between scene recognition and time discrimination for neurons that showed significant phase precession during recognition and/or time discrimination (i.e., blue plus green circles in d). Note that in (e-g), more negative correlations indicate stronger phase precession. Dashed lines indicate the example hippocampal neuron shown in (a-c).

We then asked how phase precession propensity varied across the brain. Only the hippocampus and amygdala contained a significantly higher proportion of phase precession neurons than expected by chance during all three task stages (Fig. 5a; *p* < 0.05, permutation test). While substantially rarer, the proportion of phase precession neurons was also larger than expected by chance in the parahippocampal gyrus, orbitofrontal cortex, and anterior cingulate cortex, but only during encoding and not during either of the memory retrieval tests (Fig. 5a; *p* < 0.05, permutation test).

**Figure 5.**
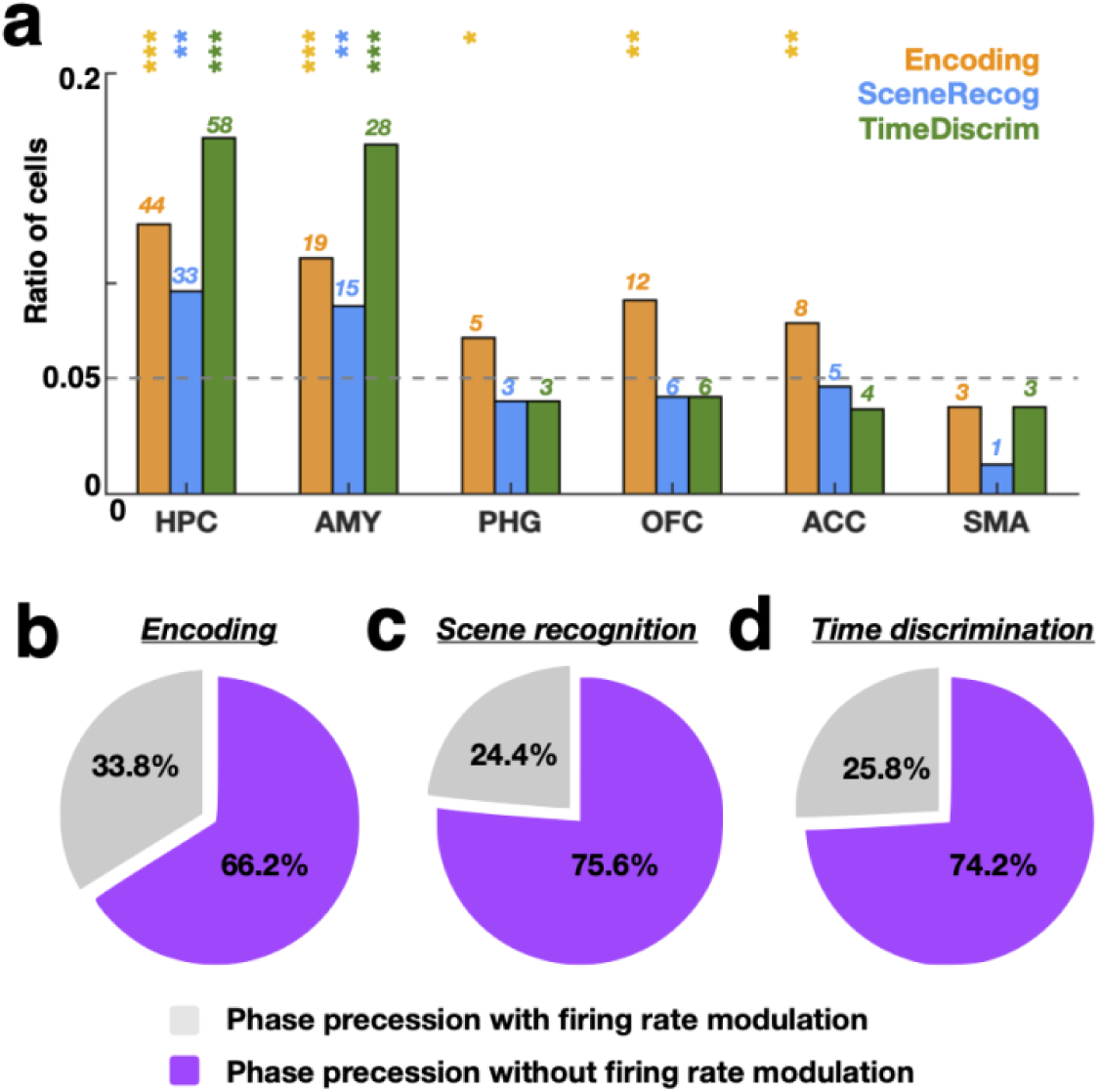
Anatomical distribution of phase precession neurons and their co-occurrence with firing rate changes. **(a)** Phase precession was most prominent in MTL. Bars demonstrate the proportion of neurons that exhibited phase precession in each anatomical area during different task stages (orange: encoding; blue: scene recognition; green: time discrimination). The number of phase precession neurons in each anatomical area is listed on the top of each bar. Dashed horizontal line indicates the chance level. Asterisks mark brain areas under specific task stage with proportions of phase precession neurons larger than expected by chance (*p* < 0.05, binomial test). HPC = hippocampus, AMY = amygdala, PHG = parahippocampal gyrus, OFC = orbitofrontal cortex, ACC = anterior cingulate, SMA = supplementary motor area. (**b**) Most phase precession neurons did not exhibit modulation by firing rate. Shown is the proportion of phase precession neurons in MTL with/without firing rate modulation (defined as firing rate differences between before vs. after boundaries during encoding). (**c** and **d**) Same as (b) but assessing whether neurons changed their firing rate when comparing before and after image onset during scene recognition (c) or time discrimination (d). \*\*\**p*< 0.001, \*\**p*< 0.01, \**p*< 0.05, n.s. = not significant.

### Distinct phase and rate coding

We previously showed that a subset of MTL neurons selectively increased their firing rates after cognitive boundaries during encoding (we labeled these cells ‘cognitive boundary cells’, Zheng et al 2022). Were the phase-precessing neurons we describe here also cognitive boundary cells? On average, phase precession neurons had higher firing rates compared to non-phase precession neurons following onset of cognitive boundaries, indicating that the two groups of neurons might overlap (Supplementary Fig. 4b-4d). However, most of the phase precession neurons (encoding: 45/68, 66.2%) were not cognitive boundary cells (Fig. 5b). Similarly, during scene recognition and time discrimination, most phase precession cells did not change their firing rate following stimulus onset relative to baseline (Fig. 5c, scene recognition: 39/51, 76.5%; Fig. 5d, time discrimination: 66/89, 74.2%).

### Phase precession strength predicts participants’ memory performance

Given the largely separate groups of cells exhibitng phase precession and firing rate changes, we asked whether phase precession and firing rate provided complimentary information about memory encoding and/or retrieval success as assessed behaviorally. We tested this idea by computing phase precession strength (correlation coefficient) and average firing rates (normalized to baseline) separately for trials with correct and incorrect memory (see *Methods*). We then quantified how well phase precession strength and/or firing rates explained participants’ memory performance (correct vs incorrect) using a generalized linear model. Based on comparisons between models with different combinations of features, we found that the winning model for explaining recognition memory performance is the model that has access to the three variables, phase precession strength during encoding, phase precession strength during retrieval, and firing rate during encoding (Fig. 6a, winning model marked in red). This model performed better than one with only access to firing rate. Similarly, for explaining time order retrieval performance, the winning model was also the one with access to all four variables (Fig. 6b; winning model marked in red). Further examining the winning models (marked in red in Fig. 6a and 6b) revealed that in both instances, phase precession strength explained most variance compared to the firing rates (Fig. 6c-d).

**Figure 6.**
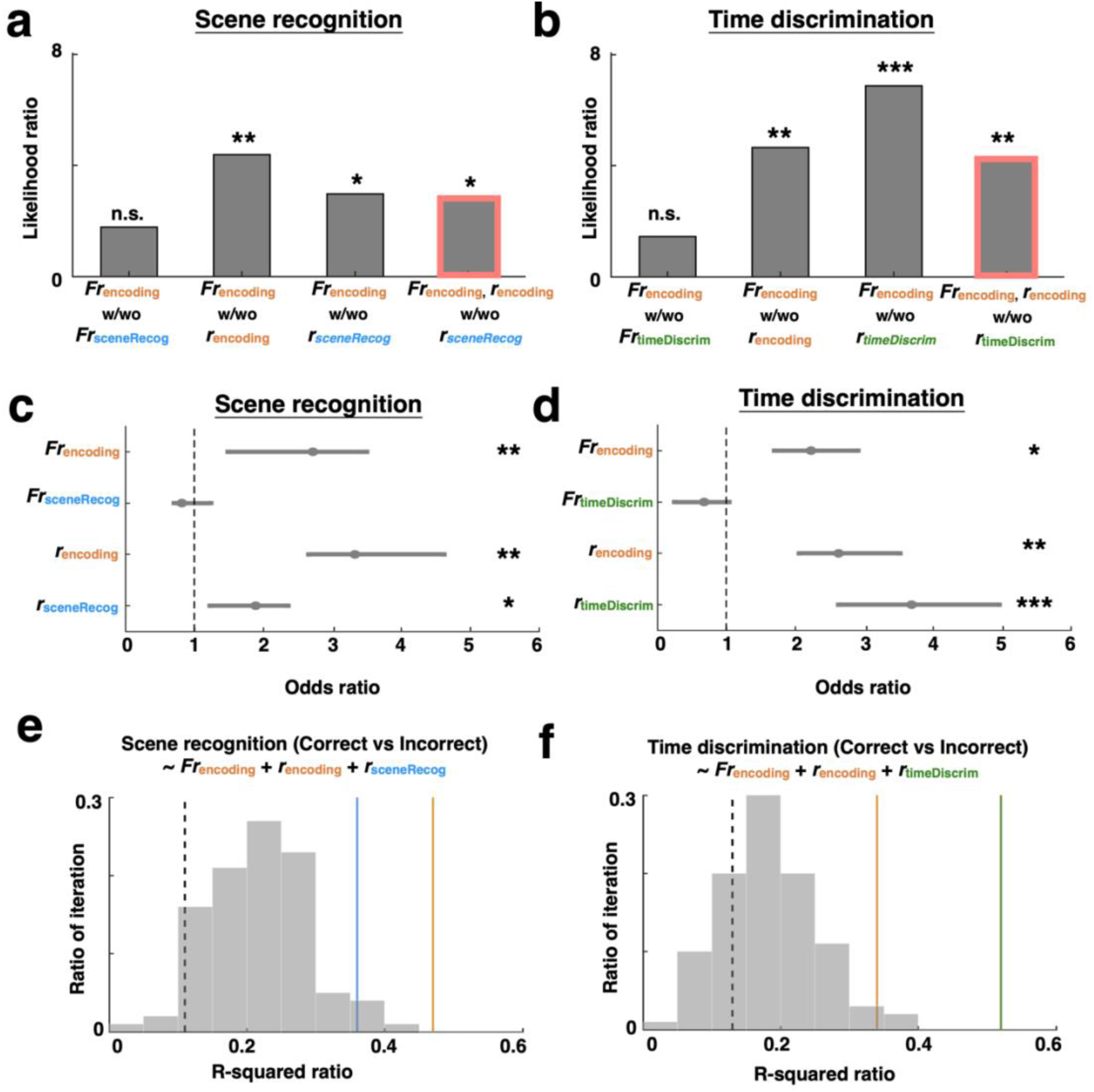
Strength of phase precession is predictive of memory encoding and retrieval success. **(a)** Model comparisons for neural activity during scene recognition task. Each bar shows a comparison between the full GLM model and reduced GLM models with/without a given predictor. Likelihood ratio bigger than 1 with significant *p*-value indicates a better model performance in explaining participants’ behavior outcomes with the added predictor. Predictors considered are firing rate during encoding (*Fr*_encoding_) and scene recognition (*Fr*_sceneRecog_), and phase-precession strength during encoding (*r*encoding) and scene recognition (*r*_sceneRecog_). (**b**) Same as (a), but for the time discrimination task. Predictors considered are firing rate during encoding (*Fr*_encoding_) and time discrimination (*Fr*_timeDiscrim_), and phase precession strength during encoding (*r*_encoding_) and time discrimination (*r*_timeDiscrim_). All the models in (a and b) consider all the recorded neurons in the medial temporal lobe. **(c and d)** Odds ratios for different predictors when predicting participants’ memory performance during scene recognition (c) and time discrimination (d). The ends of the horizontal lines indicate the confidence level and asterisk denote significance (Waldon test). (**e**) For the winning model for scene recognition (indicated by red box in panel a), the proportion of variance in the response variable (correct vs incorrect) explained by different groups of neurons. Shown are R-squared ratios of models built using all phase precession neurons during encoding (orange line), all phase precession neurons during scene recognition (blue line), and all non phase precession neurons. The total number of neurons used for the GLM model for each group are balanced by random subsampling. To do so, the non-phase precession neuron group is subsampled 100 times, each time selecting the same number of neurons as the number of phase precession neurons present during scene recognition. Dashed line indicates the chance level. (**f**) Same as €, but for time discrimination. Shown is the proportion of variance in the behaviour explained by the winning model (indicated by red box in panel b). Shown are model fit for all phase precession neurons selected during encoding (orange), all phase precession neurons selected during time discrimination (green), and all other neurons (grey, subsampled 100 times and each time with the same number of neurons as those selected during encoding). Dashed line indicates the chance level. * *p* < 0.05, ** *p* < 0.01, *** *p* < 0.001, n.s. = not significant.

Next, we assessed how much variance in the winning models was explained by subsets of the neurons (the results in above paragraph are for all recorded neurons). Compared to the rest of the MTL neurons, phase precession neurons explained significantly more variance in the behavioral accuracy during scene recognition (correct vs incorrect), with phase precession neurons during encoding and scene recognition exceeding 100% and 96% of the R-squared values achieved for random subsets of the same number of non-phase precession neurons (Fig. 6e), respectively. Similarly, phase precession neurons also explained the most variance in time discrimination accuracy, with phase precession neurons during encoding and time discrimination exceeding 97% and 100% of the R-squared values achieved for random subsets of the same number of non-phase precession neurons (Fig. 6f), respectively. In sum, the strength of phase precession provided information about memory encoding and retrieval success that was complementary to the information provided by firing rates, thereby revealing a temporal code.

## Discussion

Theta phase precession is widely observed across species and is thought to be a neural mechanism to temporally link continuous experience and encode and retrieve these experiences into and from memory^11^. However, how phase precession facilitates memory formation and retrieval remains an open question that requires more direct neural evidence. Here we test the hypothesis that phase precession supports memory formation and retrieval of naturalistic experience in humans. We found that phase precession was prominent following cognitive boundaries during movie clip watching and when participants retrieved memories of the content and temporal structure of the previously watched movie clips. The strength of phase precession was predictive of participants’ ability to encode and retrieve the memories.

Theta phase precession was prominent despite the absence of long and continuous stretches of theta activity like that observed in rodents. This observation is consistent with work in bats, where phase precession is also prominent despite theta only being present in short bouts^16^ similar to those in humans. Consistent with previous work^27, 28^, theta-frequency band LFP fluctuations in humans tend to be non-rhythmic, except for short transient theta “bouts” (Fig. 2). The frequencies of theta bouts detected on the same recording electrode can vary across a broad range (2-10Hz; Fig. 2i), which is in line with previous observations in humans^17^ and bats^16^. Theta bouts and theta phase precession are both prevalent after cognitive boundaries during encoding (Fig. 2h and Fig. 3e) and image onset during retrieval (Supplementary Fig. 2d and 2h, Fig. 3f and 3g). However, the strength of phase precession does not solely depend on theta periodicity. Illustrating this, phase precession is stronger during memory retrieval compared to encoding despite theta bouts being rarer and variable during retrieval compared to encoding (see Supplementary Fig. 2i and 2j). Previous studies have also reported phase precession when theta frequency is altered^33^, theta-modulated spiking is reduced^34–36^, or even with little periodicity of low-frequency field potentials^15, 16^. Moreover, theoretical work shows that periodicity is not necessary for phase precession^37^. Indeed, recent work suggests unique advantages of phase coding in the absence of rhythmicity, which enables multiplexing of rich information through a broad range of oscillatory frequency^38^. Our results, together with previous literature, support the idea that phase precession can serve as a flexible neural coding principle that is effective for both narrow- and broadband theta.

What triggers phase precession? We observed the strongest phase precession following cognitive boundaries during encoding and following image onset during scene recognition and time discrimination (Fig. 3c, 3f and 3g). Both cases contain sharp visual transitions, either across two different movie scenes or from a blank screen (baseline) to a movie scene. However, not all sharp visual transitions trigger phase precession, as phase precession following movie clip onsets, movie clip offsets, image offsets, and button presses was not different from that expected by chance (Fig. 3c, 3f and 3g). We therefore posit that phase precession occurs when encoding or retrieving memories across events within a continuous experience. This hypothesis is consistent with previous findings^6, 25^ that show phase precession at event transitions in discrete event sequences.

The strength of phase precession varied across the different task stages. As stated above, stronger phase precession was observed during scene recognition and time discrimination compared to encoding (Fig. 4e and 4f). Also, during recognition memory, phase precession was stronger in old compared to new trials. We posit that these differences in phase precession strength might be the result of memory-based decision making, during which the precise temporal structure of previous experiences is relevant. Supporting this hypothesis, more phase precession neurons (Fig. 4d) were found during time discrimination compared to scene recognition. Also, stronger phase precession (Fig. 4g) was more predictive of participants’ order memory accuracy compared to recognition memory accuracy (Fig. 6c and Fig. 6d). A previous study^39^ has also reported stronger phase coding (but not phase precession) when animals were instructed to make a memory-based decision compared to passively trespassing the same environment.

A potential explanation of changes in phase precession strength between tasks could be the different strategies participants used when solving the two memory tests. During time discrimination, participants needed to determine the order of the two frames shown on the screen, which requires recollection of the entire event sequence. In contrast, during scene recognition, participants only needed to retrieve the memory associated with a single image. Notably, during scene recognition, phase precession is mostly observed during trials in which an old (familiar) image was shown (Supplementary Fig. 5). Identification of novel (foil) images might not require phase precession because it could rely on novelty signals^40, 41^. An important future question is what the relationship is between phase precession and other forms of neural coding, such as novelty ^40^. The dynamic task-dependent phase precession strength observed in our study suggests that phase precession can serve different functional roles as needed.

Models with access to both phase precession strength and firing rate were able to explain variance in behaviorally assessed memory accuracy of the participants significantly better than models with access to firing rate alone (Fig. 6). This finding supports the idea that phase information provides information that trial-averaged firing rate does not provide, making the two codes complementary^42, 43^. Combining both rate and phase coding, hippocampal cell assemblies may generate sequential structures across single theta cycles that represent sequences of past, current, and future states in both the spatial and episodic domain in a compressed manner^12, 13, 44^. While phase precession has largely been reported for neurons with known tuning curves (i.e., rate coding, such as place cells and grid cells), most phase precession neurons in our study did not show modulation of firing rate with the variables examined in this study (Fig. 5b-d). This independence of phase precession strength and firing rate modulation is consistent with theoretical models^43, 45^, and previous observations in rodents^46^ and humans^17^ that show that phase precession can appear in the absence of concurrent firing rate changes. An alternative interpretation is that we did not test for the variables that would modulate firing rate in these neurons. Even for the subgroup of neurons which showed both significant firing rate changes and phase precession, our findings differed from those of place cells (which exhibit both rate and phase coding). This is because in our study, all movie clips are shown only once (they are novel). Contrary to place cells^11^, the rate tuning of phase precession neurons in our study is thus not tied to a specific stimulus. These findings suggest that phase precession in a given neuron can occur during exposure to many different novel stimuli that have the occurrence of cognitive boundaries in common, thereby flexibly capturing the temporal structure of diverse event sequences using a shared neural coding mechanism.

In sum, we demonstrate non-spatial phase precession during memory formation and retrieval of naturalistic experience in humans. The strength of phase precession and the groups of neurons showing phase precession are task specific and the strength of phase precession was predictive of the success of memory encoding and retrieval above and beyond firing rates alone. Our findings suggest that phase precession in humans is a general neural coding mechanism that can flexibly support different aspects of episodic memory.

## Methods

### Task

The task (Fig. 1a-d) consisted of three parts: encoding, scene recognition, and time discrimination. During encoding, participants watched 90 novel and silent clips embedded with or without boundaries, defined as movie cuts transitioning to scenes from the same or different movie. To assess attention, a yes/no question related to the content of the clip appeared randomly after every 4-8 clips. After watching all 90 clips, participants were instructed to take a short break, roughly 5 minutes, before proceeding to the memory tests. Participants first performed the scene recognition test and were instructed to identify extracted frame from watched clips as “old” and novel frames as “new”. Participants then performed the time discrimination test to identify whether the “Left” or “Right” frame was shown first (earlier in time) during encoding. More detailed information about the task design can be found in our previous publication ^26^.

### Participants

Twenty-two patients (13 females, mean age = 39 ± 16 years, mean ± s.t.d, see participants’ demographics in Supplementary Table 1) with refractory epilepsy volunteered for this study and provided their informed consents. Nineteen of these participants come from our previous work ^26^ and there are three new participants. Participants performed the task while they stayed at the hospital and were implanted with electrodes for seizure monitoring. The study protocol was approved by the Institutional Review Boards at Toronto Western Hospital and Cedars Sinai Medical Center. The location of the implanted electrodes was solely determined by clinical needs.

### Electrophysiological recordings

Broadband neural signals (0.1 – 8000Hz filtered) were recorded using Behnke-Fried electrodes (Fried et al., 1999) (Ad-Tech Medical, Wisconsin, USA) at 32KHz using the ATLAS system (Neuralynx Inc., Montana, USA). We recorded bilaterally from the amygdala, hippocampus, and parahippocampus, and other regions outside the medial temporal lobe. Electrode locations were determined by co-registering postoperative CT and preoperative MRI scans using Freesurfer’s mri_robust_register^47^. Each Behnke-Fried electrode shank had 8 microwires on the tip and were marked as one anatomical location. To assess potential anatomical specificity across different participants, we aligned each participant’s preoperative MRI scan to the CIT168 template brain in MNI152 coordinates ^48^ using a concatenation of an affine transformation followed by a symmetric image normalization (SyN) diffeomorphic transform^49^. The fifty electrodes within the medial temporal lobe across all the participants are illustrated on the CIT168 template brain in Fig. 1e, with corresponding MNI coordinates listed in Supplementary Table 2.

### Spike sorting and quality metrics of single units

Spike sorting was performed offline using a semi-automated template matching algorithm Osort^50^. See Zheng et al. 2022 for more details on spike sorting procedures. We identified 1103 neurons in this dataset across all brain areasconsidered. As an accurate assesment of phase precession requires a sufficient number of spikes, we excluded 207 neurons with low firing rates (< 0.5Hz) and analyzed the remaining 936 neurons (503 neurons in the medial temporal lobe). The quality of our spike sorting results was evaluated using our standard set of spike sorting quality metrices^51, 52^ for all considered 936 putative single neurons (Supplementary Fig. 1).

### Characteristics of theta oscillations

#### Pre-processing

Local field potentials (LFP) were recorded simultaneously with single neuron activity and was used for computing phase precession along with the spike activity from neurons detected from the same microwire. To eliminate potential influences of the spike waveform on the higher frequency parts of the LFP^53^, we replaced the LFP in a 3ms long time window centered on the detected spike by linear interpolation. We then downsampled this spike-free version of the LFP from 32kHz to 250Hz, followed by further post-processing using automatic artifact rejection^54^ and manual visual inspection using function *fr_databrowser.m* from Fieldtrip toolbox^55^. Trials with large transient signal changes were removed from further analyses. Examples of pre-processed LFP are shown in Fig. 2a, 2c, 2e and gray lines in Fig. 2g and 2i.

#### Spectral analysis

The power spectra (1-20Hz) of the pre-processed LFP were computed using Welch’s method (function *pwelch.m* in MATLAB) with Hamming windows (50% overlap). In Fig. 2k and 2n, we performed the spectral analysis on the pre-processed LFP within the 0 to 1 seconds time window that follows boundaries in boundary clips.

#### Theta bouts detection

To examine the cycle-to-cycle variability of theta oscillations, we detected transient theta bouts using the method described in previous papers^27, 29^. Briefly, first two versions of the LFP were computed: one low-pass filtered at 40Hz (“low-passed LFP”) for identifying peaks and troughs in a “smooth” version of LFP without high frequency components; and one band-pass filtered within the theta range (2-10Hz, “theta-filtered LFP”) to identify zero-crossings as searching windows for theta bouts detection. Theta bouts (2-10Hz) were identified as time periods in which all detected cycles had similar amplitudes (amp_consistency_threshold = 0.6), similar periods (period_consistency_threshold = 0.6), and relatively symmetric rise and decay flanks (monotonicity_threshold = 0.6) for more than three consecutive putative cycles in time. The frequency of theta bouts was computed as the ratio of how many cycles the theta bout had divided by how long in seconds it lasted. For example, a theta bout that lasted for 2 seconds with 4 cycles had a frequency of 2Hz. The time ratio of theta bouts was computed as how much time of the analysis window was occupied by theta bouts divided by the total length of the analysis window. For example, a theta bout that appeared for 1 second out of a 2-second time window had a time ratio of 0.5.

#### Spike phase estimation

Based on the peaks, troughs and zero-crossings identified as described above, we estimated theta phase at all points of time by linearly interpolating between the peaks (0° or 360°), troughs (180°) and zero-crossings (90° or 270°) cycle by cycle. Compared to a conventional Hilbert transform approach, this phase-interpolation method eliminates potential distortions introduced when estimating the phase of non-sinusoidal LFPs^27, 56^. To reduce nose in the phase estimation, which is critical for phase precession analysis, we excluded spikes that occurred during the lowest 25^th^ percentile of theta power distribution^16^. The phase assigned to a given spike was set equal to the phase estimated at the point at which the action potential was at its peak.

### Phase precession measurements

#### Spike-phase circular linear correlation method (Method 1)

We analyzed spikes that occurred within the first three theta cycles following boundaries during encoding or the onset of image display during scene recognition and time discrimination. Phase precession was quantified using circular statistics. For each neuron, we computed the circular-linear correlation coefficient ^31^ between the spike phase (circular) and time in unwrapped theta phases (linear). Unwrapped theta phase was defined as accumulated cycle-by-cycle theta phase from the alignment point, such as boundary. To assess statistical significance, we generated a null distribution for each circular-linear correlation using surrogate data generated from shuffling neurons spike timing for 1000 times. This procedure maintained the firing rate and spike phase distribution of each neuron while scrambling the correspondence between the spike phase and spike time within each trial. A neuron was considered as a significant phase precession neuron if the observed circular linear correlation exceeded the 95^th^ percentile (*p* < 0.05) of the surrogate null distribution of the correlation coefficient.

#### Chance level of neurons showing phase precession

We estimated the number of neurons exhibiting significant phase precession by chance by recomputing the spike-phase circular linear correlation 1000 times using the surrogate data generated by shuffling trial numbers between the spike phases and LFP. For each iteration, we obtained the proportion of selected phase precession cells relative to the total number of neurons within each brain region. These 1000 values formed the empirically estimated null distribution for the proportion of phase precession cells expected by chance. A brain region was considered to have a significant amount of boundary cells or event cells if its actual fraction of significant cells exceeded 95% of the null distribution (Fig. 5a; *p* < 0.05).

### Control analyses for phase precession

We performed following control analyses to further understand the observed phase precession phenomena in our study.

#### Phase precession with different theta cycle numbers

we re-assessed phase precession while varying the number of theta cycles (3 cycles *vs.* 4 cycles) from which the spikes were included for the same analyses. We found that more theta cycles resulted in more spikes included for the analyses but did not significantly affect the proportion of cells exhibiting phase precession (3 cycles *vs*. 4 cycles comparison, encoding: *χ*^2^= 5.17, *p* = 0.5, scene recognition: *x*^2^ = 7.32, *p* = 0.25, time discrimination: *x*^2^ = 6.22, *p* = 0.3, chi-square test).

#### Phase precession within various analyses windows

we evaluated the prevalence of phase precession by computing the spike-phase circular linear correlation at various task windows (Fig. 3e-g). For example, first three theta cycles after 4-second of no boundary clips, baseline, clip onsets and clip offsets in all the clips, and image offsets during scene recognition and time discrimination.

#### Phase precession using spike-phase autocorrelation method (Method 2)

To examine the robustness of the observed phase precession, we employed a second method to further estimate phase precession based on comparing the autocorrelation of spiking of each neuron with the frequency of the underlying LFP ^32^. This alternative method is widely used in rodents^57, 58^ with stereotypical 8Hz theta, and also in animals with nonstationary theta, such as bats^16^, non-human primates^59–61^ and humans^17^. For each neuron, we assigned to each spike its unwrapped theta phase by accumulating the cycle-by-cycle phase following boundary during encoding. We computed the autocorrelation (‘phase autocorrelation’) of these cumulative spiking phases of the spike-train, using 60° bins with window length of 3 cycles. If a neuron exhibits phase precession, its firing phases will occur more and more ahead of theta cycles ^32^. In other words, its spike phase autoccorelation will have peaks at a higher rate than the theta cycles (Supplementary Fig. 3f and 3h). To measure this, we fitted decaying sine wave functions to the spike phase autocorrelation for visualization and computed the power spectra of the fitted lines using Fourier transform. The result was presented as a power spectral density plot of the relative frequency ratio between the spiking frequency and the theta frequency. Based on the obtained power spectrum density plot (insets in Supplementary Fig. 3e-3h), we computed the modulation index^17^, defined as the amplitude of the power spectra peak normalized by the total amplitude across all other relative frequencies. We then generated a null distribution of modulation indices from spike phase autocorrelations based on surrogate data generated using the same shuffling procedure discussed in spike-phase circular linear correlation. A neuron was considered as exhibiting significant phase precession if its observed modulation index exceeded the 95^th^ percentile of the surrogate distribution. To eliminate potential bias from low spike counts, we only analyzed neurons with more than 80 spikes within the analysis windows across trials for computing spike phase autocorrelation. Further, neurons showing significant phase-locking (*p* < 0.05, Rayleigh test) and a peak relative frequency around 1 (0.95 to 1.05) were excluded to ensure that we did not mistakenly identify phase locking neurons as exhibiting phase precession. Comparing the neurons selected as phase precessing between Method 1&2 shows significant agreement in which neurons are selected. For example, during encoding, 70% of the neurons selected as phase precessing with Method 1 were also selected with Method 2 (48/68, 70.5%; see Supplementary Table 3).

#### Phase precession using absolute time versus time in unwrapped theta phases

we recomputed phase precession using time in second instead of time in unwrapped theta phases for both Method 1 and Method 2 (Supplementary Fig. 3e and 3g). We recomputed the circular-linear correlation and modulation index from spike-time autocorrelation histogram within the 0 to 1 second time window after boundaries during encoding and after the onsets of image display during scene recognition and time discrimination. The alternative measurement using time in seconds resulted in the identification of a smaller number of phase precession cells as compared to our analyses using time in unwrapped theta phase (spike-time correlation *vs* spike-phase correlation: n = 22 *vs* 68, *x*^2^ = 16.6, *P* < 3x10^-4^, chi-square test; spike-time autocorrelation *vs* spike-phase autocorrelation: n = 17 *vs* 51, *x*^2^ = 14.7, *P* < 7 x 10^-3^, chi-square test). The differences of correlation coefficients when computing using time in unwrapped theta phases versus using time in seconds increased along with the variance of theta bouts’ frequency across trials (Supplementary Fig. 3k and 3l). In other words, the approach using time in unwrapped theta phases was sensitive and appropriate for capturing phase precession when LFPs oscillated at a broad theta frequency range across trials.

### Firing rate modulation

For neurons showing significant phase precession during encoding and retrieval, we computed their average firing rates within the same analysis time windows for computing phase precession - three cycles after boundaries during encoding and after image onsets after scene recognition and time discrimination. A neuron was considered having firing rate modulation if its average firing rate significantly differed between post versus pre analysis time windows (*p* < 0.05, paired two-tailed *t*-test). We also assessed the overlap between neurons showing firing rate modulation and phase precession at different task stages (Fig. 5b-d).

### Relationship between phase precession and memory performance

#### Generalized linear model

We assessed the relationship between neural metrics (firing rate, phase precession) and participants’ memory as assessed behaviorally using a generalized linear model (GLM). Using scene recognition as an example, we grouped trials into the two categories “correctly recognized” and “incorrectly recognized” depending on the accuracy of their response. To account for the difference in trial numbers, we subsampled correct trials with the same trial number as incorrect trials. We first computed phase precession (spike-phase circular linear correlation) and average firing rates (z-scored normalized to the baseline) within three theta cycles relative to cognitive boundaries during encoding and relative to image onsets during scene recognition, separately for correct and incorrect trials. For each neuron, this resulted in four values each for correct and incorrect trials: correlation coefficients specifying phase precession strength (*r_encoding_, r*_sceneRecog_) and normalized firing rates (*Fr_encoding_, Fr*_sceneRecog_). We used a mixed-effect GLM using the function *fitglme.m* from MATLAB with logistic regression and Bernoulli distribution to predict participants’ binary memory outcomes (i.e., correct vs incorrect) during scene recognition. The fixed effects were different combinations of the four values (*r_encoding_, r*_sceneRecog_, *Fr_encoding_, Fr*_sceneRecog_) from all the neurons in the medial temporal lobe. The random effects were neuron ID nested within session ID.

#### Contribution of different fixed effects

To assess the extent to which firing rates and phase precession predicted behavior, we compared different reduced versions of above GLM model. We compared model 1 (fixed effect: *Fr_encoding_*) with model 2 (fixed effect: model 1 + *Fr*_sceneRecog_), model 3 (fixed effect: model 1 *+ r_encoding_*), and model 4 (fixed effect: model 1 *+ r_sceneRecog_*). We also added the effect of phase precession during memory retrieval (*r_sceneRecog_*) to model 3 and compared the model performance between the two. Model comparisons were performed based on the Akaike information criteria, expressed as a log likelihood ratio in Fig. 6a. We did this separately also for the time discrimination task, in which all variables were replaced by the equivalent for the time discrimination task (Fig. 6b). We quantified the strength of each fixed effect using odds ratios to test if they have strong relationship with participants’ correct responses during scene recognition and time discrimination. For neurons (n = 77) showing phase precession during encoding or/and scene recognition, we evaluated whether they show significant phase precession or firing rate modulation for trials they correctly versus failed to recognize tested images. An odds ratio is defined as follow.

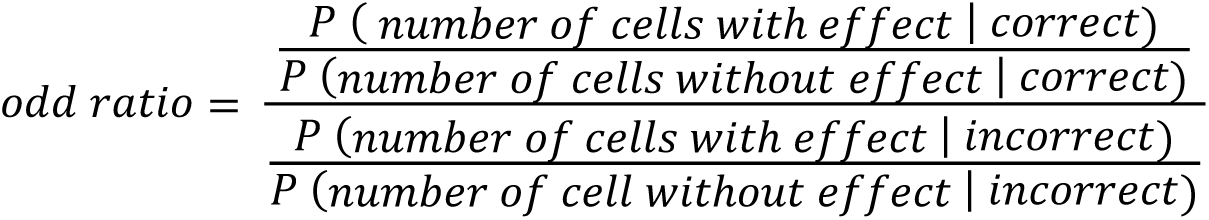

Effect here refers to one of the following: significant firing rate modulation during encoding (*Fr_encoding_*) or scene recognition (*Fr*_sceneRecog_), significant phase precession during encoding (*r_encoding_*) or scene recognition (*r*_sceneRecog_). We then performed a Wald test (function waldtest.m from MATLAB) to access the significance of computed odds ratio. As shown in Fig.6c, significant odds ratio bigger than 1 indicated that participants more likely got correct when the given effect was present. We applied same methods for time discrimination as well (Fig. 6d).

#### Contribution of different cell groups

We then assessed the ability of different groups of neurons in predicting participants’ memory outcomes (correct vs incorrect). To do so we fit the same GLM model 4 as described above (fixed effects: *r_encoding_ + Fr_encoding_ + r*_sceneRecog_ or *r*_timeDiscrim_) but including only the neurons who showed phase precession during encoding, only neurons showing phase precession during scene recognition or time discrimination, and the non-phase precession neurons. We then computed the R-squared ratio as the ratio between the R-squared value of GLM Model 4 using different subgroups of neurons 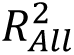 versus all medial temporal lobe neurons 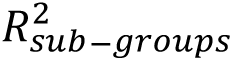, as follow:

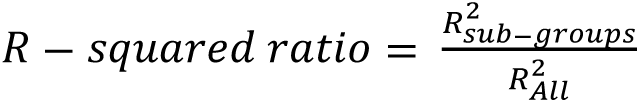

We subsampled the non-phase precession neurons to balance the number of neurons across different subgroups when building the GLM models.

### Statistical methods and software

Participants were not informed of the existence of cognitive boundaries in the clips. All the statistical analyses were conducted in MATLAB, primarily using the Statistics and Machine Learning toolbox. For comparison against specific values, we used one-sample *t*-test. For comparison between two groups, we primarily used the Kolmogorov–Smirnov test while for omnibus testing, we used ANOVA, unless otherwise specified in the text. When the normality of dataset was not clear, non-parametric permutation tests were used to determine the significance level by comparing the real test-statistic to the null distribution estimated from surrogate dataset.

---

**Supplementary Table 1.**
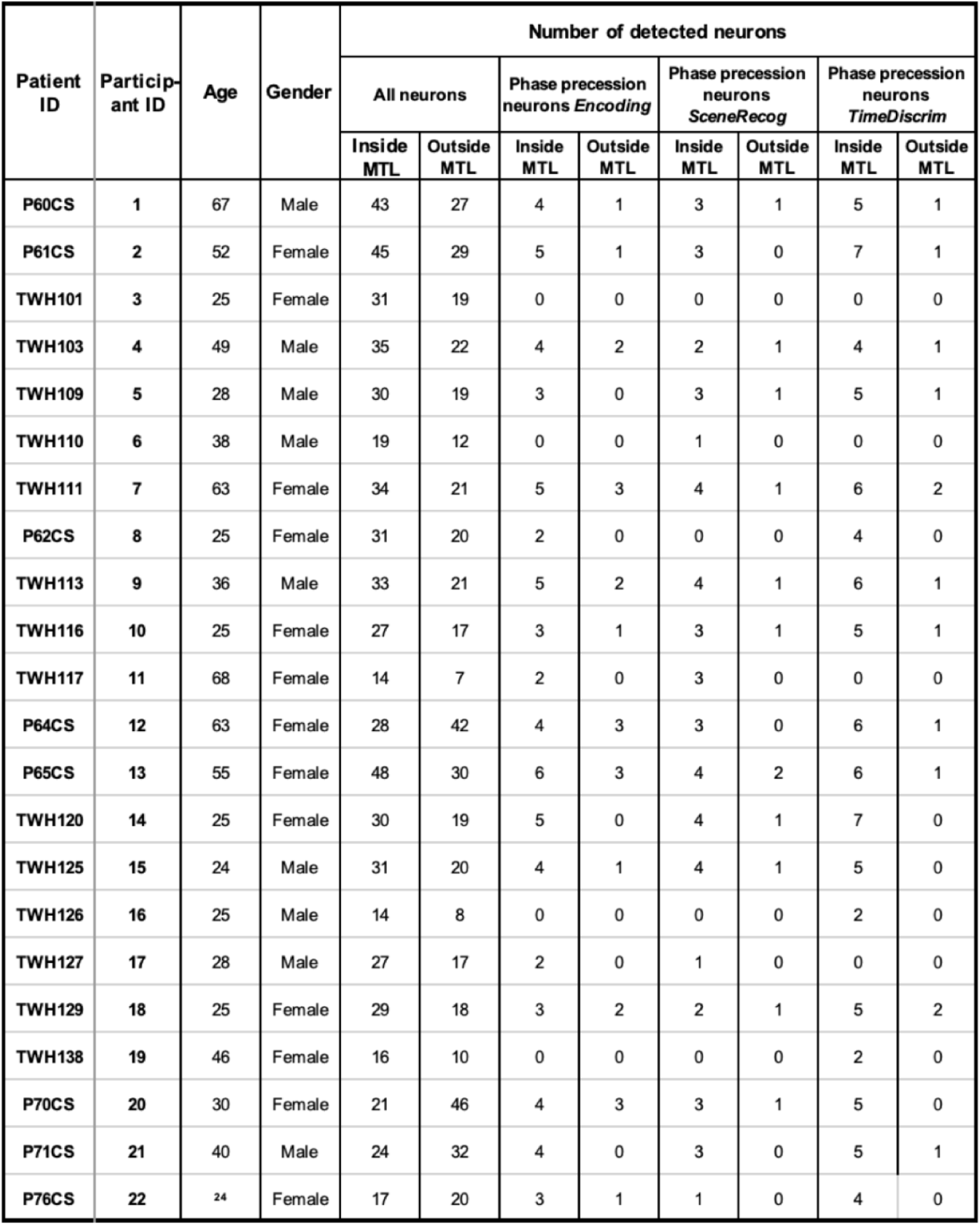
Participants’ demographics.

| Patient ID | Participant ID | Age | Gender | Number of detected neurons |  |  |  |  |  |  |  |
| --- | --- | --- | --- | --- | --- | --- | --- | --- | --- | --- | --- |
|  |  |  |  | All neurons |  | Phase precession neurons *Encoding* |  | Phase precession neurons *SceneRecog* |  | Phase precession neurons *TimeDiscrim* |  |
|  |  |  |  | Inside MTL | Outside MTL | Inside MTL | Outside MTL | Inside MTL | Outside MTL | Inside MTL | Outside MTL |
| P60CS | 1 | 67 | Male | 43 | 27 | 4 | 1 | 3 | 1 | 5 | 1 |
| P61CS | 2 | 52 | Female | 45 | 29 | 5 | 1 | 3 | 0 | 7 | 1 |
| TWH101 | 3 | 25 | Female | 31 | 19 | 0 | 0 | 0 | 0 | 0 | 0 |
| TWH103 | 4 | 49 | Male | 35 | 22 | 4 | 2 | 2 | 1 | 4 | 1 |
| TWH109 | 5 | 28 | Male | 30 | 19 | 3 | 0 | 3 | 1 | 5 | 1 |
| TWH110 | 6 | 38 | Male | 19 | 12 | 0 | 0 | 1 | 0 | 0 | 0 |
| TWH111 | 7 | 63 | Female | 34 | 21 | 5 | 3 | 4 | 1 | 6 | 2 |
| P62CS | 8 | 25 | Female | 31 | 20 | 2 | 0 | 0 | 0 | 4 | 0 |
| TWH113 | 9 | 36 | Male | 33 | 21 | 5 | 2 | 4 | 1 | 6 | 1 |
| TWH116 | 10 | 25 | Female | 27 | 17 | 3 | 1 | 3 | 1 | 5 | 1 |
| TWH117 | 11 | 68 | Female | 14 | 7 | 2 | 0 | 3 | 0 | 0 | 0 |
| P64CS | 12 | 63 | Female | 28 | 42 | 4 | 3 | 3 | 0 | 6 | 1 |
| P65CS | 13 | 55 | Female | 48 | 30 | 6 | 3 | 4 | 2 | 6 | 1 |
| TWH120 | 14 | 25 | Female | 30 | 19 | 5 | 0 | 4 | 1 | 7 | 0 |
| TWH125 | 15 | 24 | Male | 31 | 20 | 4 | 1 | 4 | 1 | 5 | 0 |
| TWH126 | 16 | 25 | Male | 14 | 8 | 0 | 0 | 0 | 0 | 2 | 0 |
| TWH127 | 17 | 28 | Male | 27 | 17 | 2 | 0 | 1 | 0 | 0 | 0 |
| TWH129 | 18 | 25 | Female | 29 | 18 | 3 | 2 | 2 | 1 | 5 | 2 |
| TWH138 | 19 | 46 | Female | 16 | 10 | 0 | 0 | 0 | 0 | 2 | 0 |
| P70CS | 20 | 30 | Female | 21 | 46 | 4 | 3 | 3 | 1 | 5 | 0 |
| P71CS | 21 | 40 | Male | 24 | 32 | 4 | 0 | 3 | 0 | 5 | 1 |
| P76CS | 22 | 24 | Female | 17 | 20 | 3 | 1 | 1 | 0 | 4 | 0 |

**Supplementary Table 2.** Electrode locations. MNI coordinates of microwire bundles in the medial temporal lobe.

| Electrode ID | PatientID | Location | Side | MNI coordinates |  |  |
| --- | --- | --- | --- | --- | --- | --- |
|  |  |  |  | X | Y | Z |
| 1 | TWH101 | Hippocampus | Left | -20.98 | -23.8 | -13.93 |
| 2 | P64CS | Hippocampus | Left | -20.44 | -17.61 | -11.61 |
| 3 | P60CS | Hippocampus | Left | -27.09 | -16.5 | -14.95 |
| 4 | TWH109 | Hippocampus | Left | -26.63 | -21.1 | -16.32 |
| 5 | TWH110 | Hippocampus | Left | -24.98 | -17 | -15.19 |
| 6 | TWH111 | Hippocampus | Left | -27.58 | -22.29 | -15.66 |
| 7 | TWH113 | Hippocampus | Left | -20.98 | -25.81 | -16.33 |
| 8 | TWH116 | Hippocampus | Left | -24.22 | -18.17 | -18.11 |
| 9 | TWH125 | Hippocampus | Left | -22 | -22.81 | -20.9 |
| 10 | TWH129 | Hippocampus | Left | -23.11 | -18.17 | -15.77 |
| 11 | P70CS | Hippocampus | Left | -20.42 | -24.85 | -12.02 |
| 12 | P71CS | Hippocampus | Left | -21.97 | -17.84 | -5.84 |
| 13 | P76CS | Hippocampus | Left | -23.35 | -19.58 | -19.57 |
| 14 | TWH103 | Hippocampus | Right | 26.63 | -15.83 | -16.49 |
| 15 | TWH109 | Hippocampus | Right | 24.8 | -11.73 | -15.77 |
| 16 | P61CS | Hippocampus | Right | 35.99 | -19.49 | -17.69 |
| 17 | P62CS | Hippocampus | Right | 32.05 | -26.82 | -6.36 |
| 18 | TWH116 | Hippocampus | Right | 24.22 | -15.24 | -19.28 |
| 19 | TWH120 | Hippocampus | Right | 29.38 | -14.6 | -22.86 |
| 20 | TWH129 | Hippocampus | Right | 26.62 | -11.81 | -20.9 |
| 21 | TWH138 | Hippocampus | Right | 31.88 | -20.64 | -14.63 |
| 22 | P70CS | Hippocampus | Right | 32.21 | -21.39 | -10.69 |
| 23 | P71CS | Hippocampus | Right | 22.12 | -15.1 | -13.93 |
| 24 | P76CS | Hippocampus | Right | 23.12 | -20.06 | -21.7 |
| 25 | TWH111 | Amygdala | Left | -22.63 | -2.78 | -19.59 |
| 26 | P64CS | Amygdala | Left | -19.09 | -4.89 | -17.3 |
| 27 | P65CS | Amygdala | Left | -18 | -3.08 | -15.11 |
| 28 | TWH113 | Amygdala | Left | -20.18 | -7.39 | -14.55 |
| 29 | TWH125 | Amygdala | Left | -23.87 | -5.52 | -24.23 |
| 30 | TWH127 | Amygdala | Left | -20.07 | -6.35 | -18.93 |
| 31 | P70CS | Amygdala | Left | -13.2 | -8.71 | -18.01 |
| 32 | P71CS | Amygdala | Left | -13.76 | -5.96 | -12.93 |
| 33 | P76CS | Amygdala | Left | -21.55 | -4.58 | -17.14 |
| 34 | P60CS | Amygdala | Right | 21.89 | -2.1 | -20.98 |
| 35 | TWH109 | Amygdala | Right | 19.93 | -3.77 | -16.14 |
| 36 | TWH113 | Amygdala | Right | 23.32 | -4.19 | -17.92 |
| 37 | TWH120 | Amygdala | Right | 21.78 | -4.59 | -24.17 |
| 38 | TWH127 | Amygdala | Right | 21.46 | -3.92 | -18.43 |
| 39 | P70CS | Amygdala | Right | 25.82 | -9.78 | -18.28 |
| 40 | P71CS | Amygdala | Right | 14.52 | -2.91 | -10.85 |
| 41 | P76CS | Amygdala | Right | 20.99 | -4.05 | -18 |
| 42 | P61CS | Parahippocampus | Left | -28.26 | -34.91 | -8.78 |
| 43 | TWH117 | Parahippocampus | Left | -17.65 | -29.59 | -15.66 |
| 44 | TWH120 | Parahippocampus | Left | -21.38 | -30.21 | -18.28 |
| 45 | TWH125 | Parahippocampus | Left | -18.57 | -38.31 | -16.98 |
| 46 | TWH127 | Parahippocampus | Left | -22 | -27.85 | -21.33 |
| 47 | TWH129 | Parahippocampus | Left | -17.95 | -33.95 | -6.57 |
| 48 | TWH125 | Parahippocampus | Right | 23.89 | -28.71 | -21.82 |
| 49 | TWH126 | Parahippocampus | Right | 17.41 | -28.87 | -10.16 |
| 50 | TWH127 | Parahippocampus | Right | 17.94 | -29.65 | -9.66 |

**Supplementary Table 3.** Method comparison. Number of phase precession neurons detected during different task stages using the two different methods spike-phase correlation and spike-phase autocorrelation. The bottom row shows the agreement between the two methods (neurons selected by both).

|  | Encoding | Scene recognition | Time discrimination |
| --- | --- | --- | --- |
| **Method 1: Circular linear correlation** | 68 | 51 | 89 |
| **Method 2: Spike-phase autocorrelation** | 51 | 42 | 73 |
| **Overlap between both methods** | 48/68<br>(70.5%) | 36/51<br>(70.6%) | 68/89<br>(76.4%) |

**Supplementary Fig. 1.**
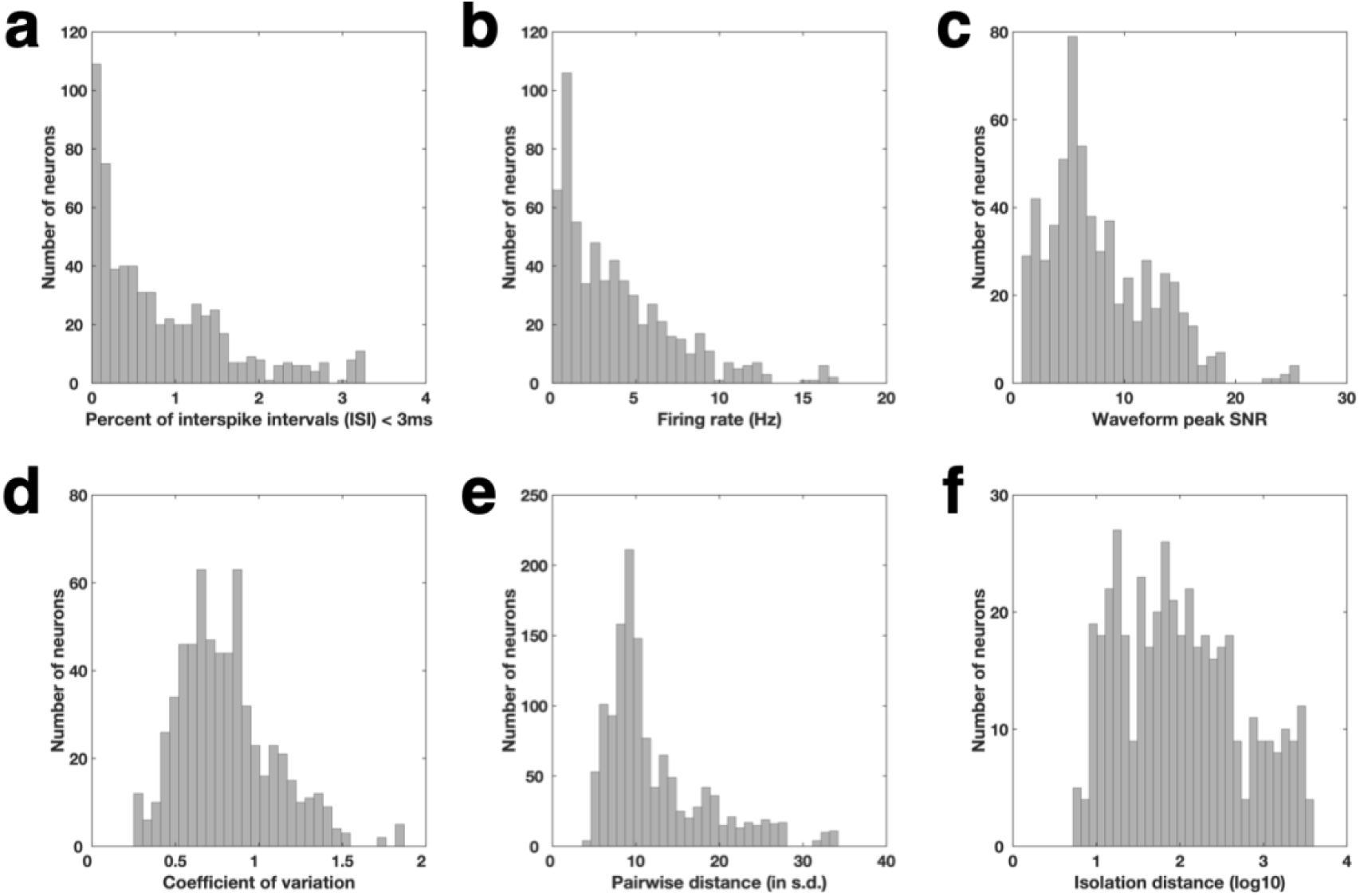
Spike sorting quality metrics for all identified putative single cells with firing rates bigger than 0.5 Hz. (a) Proportion of inter-spike intervals (ISI) that were shorter than 3ms. (b) Average firing rate within the entire recording session for all identified putative single cells. (c) Waveform peak signal-to-noise ratio (SNR), which is the ratio between the peak amplitude of the mean waveform and the s.d. of the noise of each identified putative single cell (8.00 ± 4,73, mean ± s.d.). (d) Coefficient-of-variation (CV2) in the ISI for each identified putative single cell (0.74 ± 0.24, mean ± s.d.). (e) Pairwise isolation distance between putative single cells identified from the same wire. (f) Isolation distance across all identified putative single cells that was calculated in a ten-dimensional feature space of the energy normalized waveforms.

**Supplementary Fig. 2.**
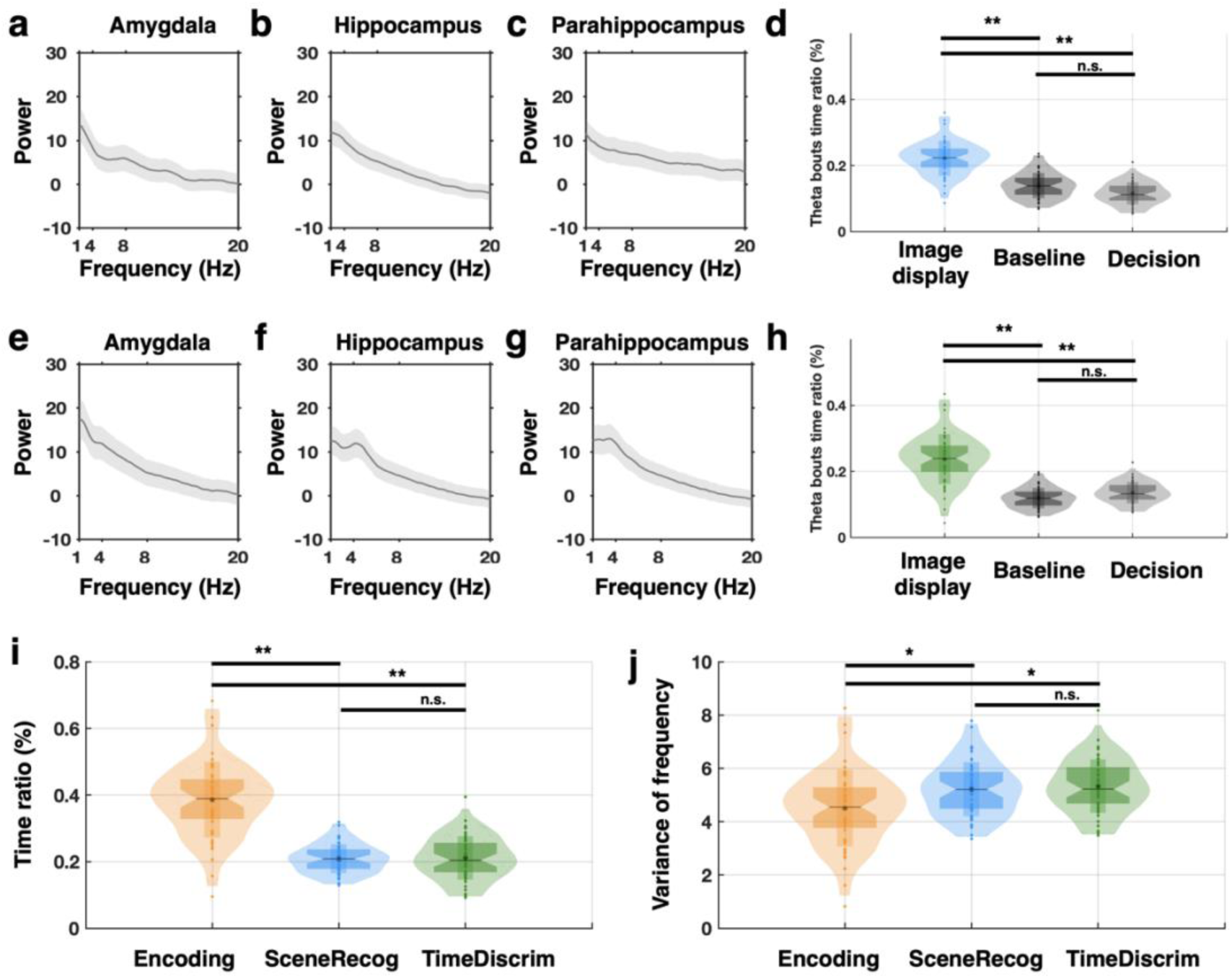
Comparison of theta bout properties between task stages. **(a**-**c)** Power spectra of local field potentials recorded during scene recognition averaged across all microelectrodes and the entire task within the indicated brain area. The shaded area indicates the standard error mean. (e-g) Same, but for time discrimination. **(d** and **h)** Proportion of the 1s long analysis window occupied by theta bouts during following the onset of image, baseline, and probe (decision) during scene recognition (d) and time discrimination (h). **(i)** Proportion of time that theta bouts occupy 0 to 1-second after cognitive boundaries (encoding) or after image display (sceneRecog and timeDiscrim). (**j**) Comparison of the variance of frequency for theta bouts that are detected within the 0 to 1-second after cognitive boundaries (encoding) or after image display (sceneRecog and timeDiscrim). * *p* < 0.05, ** *p* < 0.01, *** *p* < 0.001, n.s. = not significant.

**Supplementary Fig. 3.**
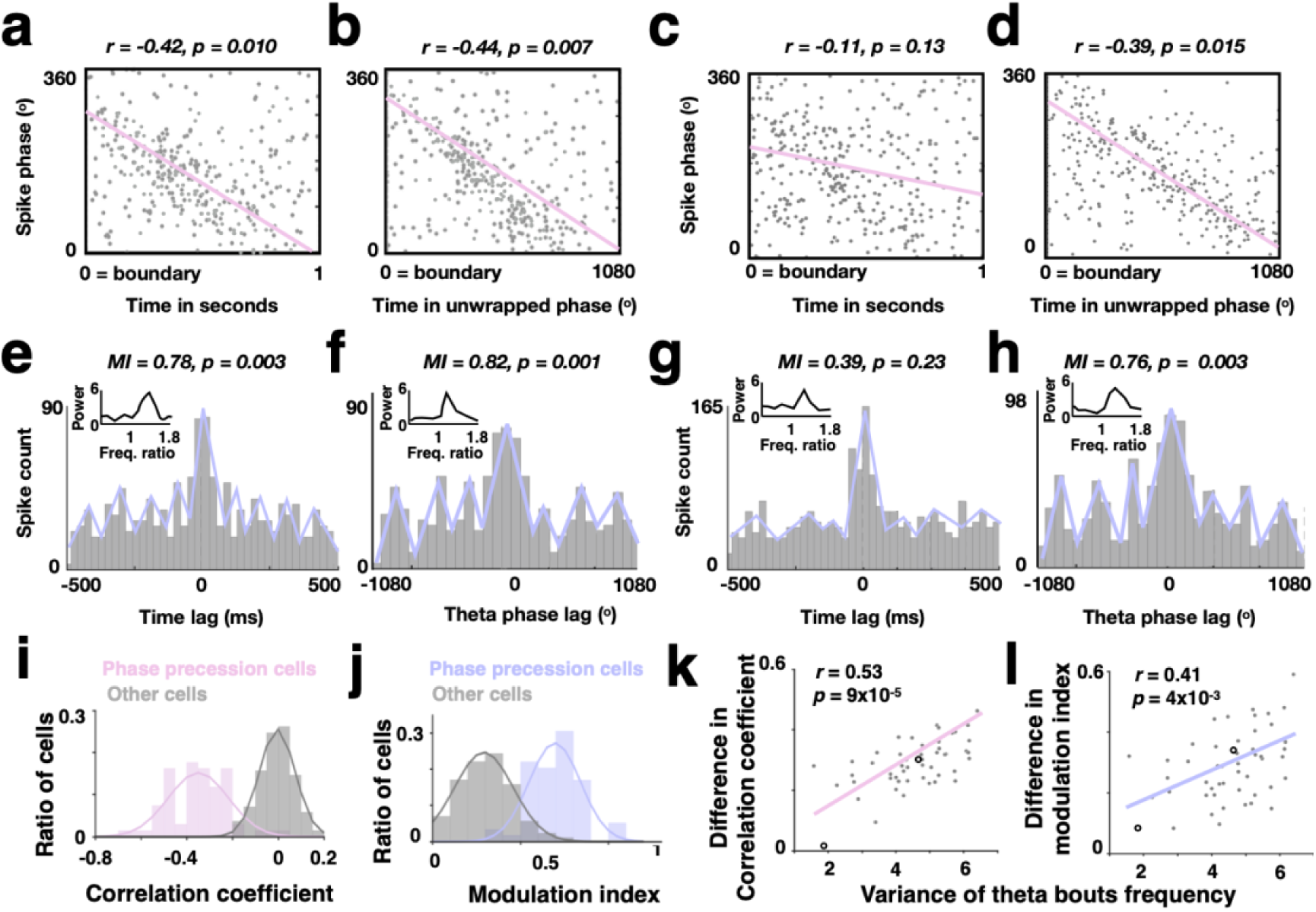
Phase precession quantified using the method 1 and 2. **(a**-**d)** Two example hippocampal neurons’ (recorded on the microelectrodes shown in Fig. 2g and 2i). Spike phases (relative to theta oscillations) are plotted as a function of time in seconds (a and c; method 1) or time in unwrapped theta phases (b and d; method 2), aligned to cognitive boundaries. Phase precession is quantified as the circular-linear correlations between neuronal spiking phases and time in seconds (a and c; method 1) or time in unwrapped theta phases (b and d; method 2). Pink lines indicate the correlation, with the correlation value and its statistical significance listed on the top of each subplot. **(e**-**h)** Further calculations for Method 2. The spike-time autocorrelation (e and g) and spike-phase autocorrelation (f and h) plots for the same neurons in (a-d). Blue lines indicate the decaying sine wave function fitted to the autocorrelation plots. Inset shows the spike-phase spectra, with power as the y-axis and the relative frequency between the frequency of spiking and frequency of heta oscillations as the x-axis. Phase precession is quantified using the modulation index (MI), which is defined as the fraction between the peak height divided by the area under the curve in the spike-phase spectra plot. The modulation index and its statistical significance are listed on the top of each subplot. (**i** and **j**) Correlation coefficients (i) and modulation indices (j) for significant (pink) versus non-significant (gray) phase precession neurons. (**k**) Differences in correlation coefficient (e.g., |*r* in b – *r* in a|) against the frequency variance of theta bouts detected in these microelectrodes for significant neurons during encoding. (**l**) Differences in modulation index (l, e.g., |*MI* in f – *MI* in e|) against the frequency variance of theta bouts detected in these microelectrodes for significant neurons during encoding. In k and l, each dot represents one neuron. The empty circle represents the two example neurons in (a-h). Color lines indicate the fitted linear regression, with the correlation value and its statistical significance listed on the top of each plot.

**Supplementary Fig. 4.**
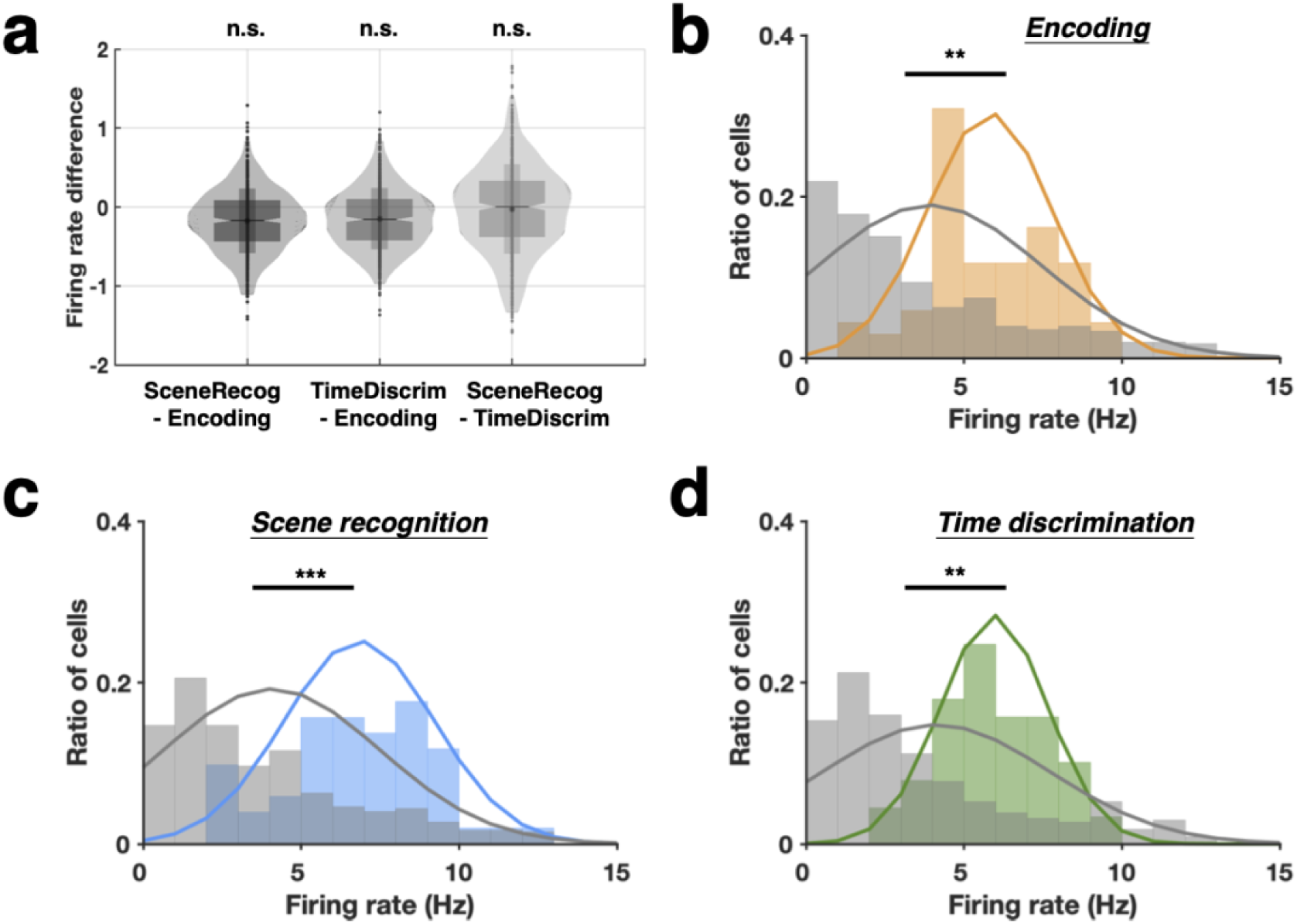
Firing rates of phase precession neurons. **(a)** For phase precession neurons (n = 117), their firing rates averaged within the 0 to 1-second time window after cognitive boundaries (encoding) and image display (scene recognition and time discrimination) are compared across different task stages. **(b-d)** Firing rates of neurons with (color) and without phase precession (gray) averaged within 0 to 1-second time window after cognitive boundaries during encoding (b), image display during scene recognition (c), and image display during time discrimination (d).

**Supplementary Fig. 5.**
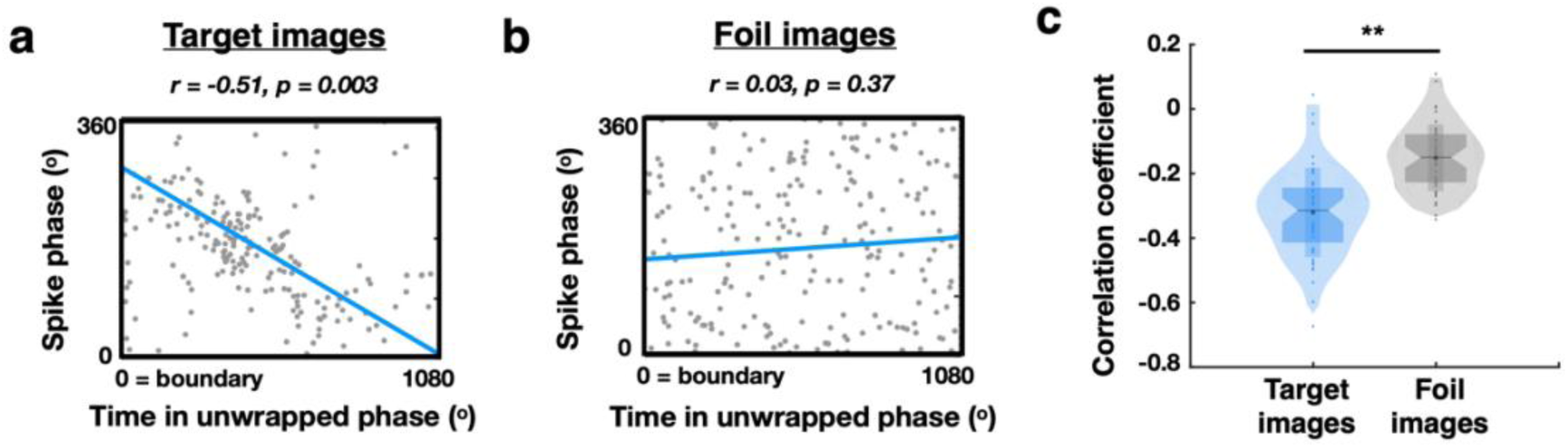
Comparison of phase precession when recognizing target versus foil images. **(a** and **b)** An example MTL neuron showing phase precession when the participant is asked to recall recognize target images (a) while no phase precession is observed when identifying foil images (b). (c) Comparison between phase precession strength (correlation coefficient values) when participants are instructed to recognize target (blue) versus foil images (gray) across all phase precession neurons identified during scene recognition (n = 51). Each dot represents one neuron. The shaded violin shape represents the data distribution with lower end of 1^st^ percentile and top end of 99^th^ percentile. The top edge and bottom edge of the shaded rectangle represent the mean ± std., respectively. \*\**p*< 0.01, n.s. = not significant.

